# Experimental Data Driven AI Framework for Flexible Protein Conformational Reconstruction

**DOI:** 10.64898/2026.03.12.708611

**Authors:** Feng Yu, Stephanie Prince, Andrew Tritt, Kanupriya Pande, Greg L. Hura, Oliver Ruebel, Susan E. Tsutakawa

## Abstract

Deep learning has revolutionized structural biology by prediction with near experimental accuracy static protein folds from amino acid sequence alone. However, proteins function as dynamic ensembles of protein conformation states, and current sequence-only models often fail to capture the specific conformational states and heterogeneity dictated by cellular environments or ligand binding. While recent generative models can sample broad conformational landscapes, they remain unconstrained by physical reality, often hallucinating plausible but experimentally invalid states. Here, we present AlphaSAXS, an end-to-end framework that constrains artificial intelligence (AI) inference using Small Angle X-ray Scattering (SAXS) experimental solution scattering data. By integrating real-space pair distance distributions (*P*(*r*)) directly into the AlphaFold architecture, AlphaSAXS effectively steers the structural hypothesis toward the experimentally observed structures. We demonstrate that AlphaSAXS resolves documented failure modes of sequence-only models in Apo-Holo transitions, successfully distinguishing between states with identical sequences but distinct scattering profiles. Furthermore, we introduce a hybrid inference protocol that couples deep learning with biophysical hydration modeling, enabling the reconstruction of solution state protein ensembles compatible with experimental data. This work establishes a paradigm for experimentally guided AI, bridging the gap between probabilistic sampling and biophysical measurement.

## Introduction

Despite the transformative achievements of AlphaFold^1^ in predicting static protein folds, the modeling of proteins with multiple conformational states remains a significant frontier in structural biology. These dynamic systems are central to vital biological processes. For example, over 40% of the human proteome consists of intrinsically disordered regions (IDRs) that exist in a dynamic conformational ensemble rather than possess a single stable structure. These ensembles are essential for many protein functions in transcription regulation,^2^ apoptosis,^3^ DNA repair,^4,5^ and liquid-liquid phase separation.^6^ Furthermore, many flexible proteins undergo critical conformational transitions upon binding to ligands or small-molecule drug interactions that are fundamental to signal transduction and modern drug discovery.^7^ A notable example is the dynamic nature of antibody complementarity-determining region (CDR) loops, where the ability to accurately sample conformational space is the prerequisite for effective paratope design.^8,9^

Current AI-based structure predictors, however, exhibit two pervasive biases that limit their ability to accurately sample conformational space. First, models based on the AlphaFold1 and AlphaFold2 architectures rely heavily on training regimes biased toward single-state representations, primarily sourced from static X-ray crystallography data. Even when incorporating NMR-derived structural ensembles from the Protein Data Bank (PDB), these models typically utilize only the first frame, enforcing a rigid “one sequence, one structure” paradigm. Second, recent diffusion-based models, such as AlphaFold3,^10^ remain poorly calibrated for IDRs. As reported by its developers, AlphaFold3 tends to generate overly compact and less accurate conformations for flexible regions, necessitating training corrections using AlphaFold2-derived data. This limitation likely stems from severe data scarcity: while the PDB contains over 200,000 structures of well-folded proteins, only a small fraction (∼10,000) are NMR-derived ensembles that represent dynamic landscapes.^11,12^ Consequently, sequence-only prediction of dynamic ensembles remains fundamentally constrained, as exemplified by the reduced accuracy of AlphaFold3 in modeling IDRs despite its advanced model architecture and training process.

To overcome these data-driven constraints, we propose a strategy that guides AI inference using direct experimental evidence. The Small-Angle X-ray Scattering (SAXS) experiment is a uniquely accessible solution-state technique that provides electron distance information suitable for acting as a structural constraint.^13,14^ SAXS offers several key advantages: it requires minimal sample preparation, has low protein requirements (∼250 µg), allows observation of protein structures in solution, and is applicable to proteins across a vast molecular weight range. Furthermore, SAXS profiles reflect the global distribution of distances between all electron pairs in a protein. In real space, this is represented by the pair distance distribution function, *P*(*r*), which provides a “molecular fingerprint” of the protein’s conformations in solution. To collect the data, X-ray scattering intensity from proteins is measured relative to their corresponding buffer as a function of the scattering angle (Fig. 1A). The buffer data is subtracted from the protein SAXS data. SAXS curves are typically presented in reciprocal space, plotting intensity versus the scattering angle, which is more precisely defined by the vector: *q* = (4π/*λ*) sin *θ*, where *λ* is the X-ray wavelength and 2θ is the scattering angle.^15^ The real space electron pair distribution, *P*(*r*), is then obtained by applying a Fourier transform.^16^ Traditional rigid-body modeling and normal mode analysis frequently distort local geometry to accommodate global scattering profiles. In contrast, integrating SAXS profiles into a deep learning framework directly guides protein structure reconstruction, minimizing modeling bias and structural violations. Once SAXS info defines the global shape of protein structures, much like a multiple sequence alignment (MSA), we hypothesized that the algorithm’s learned structural relationships would yield finer folding constraints, akin to those seen in crystallography.^17^

**Figure 1.**
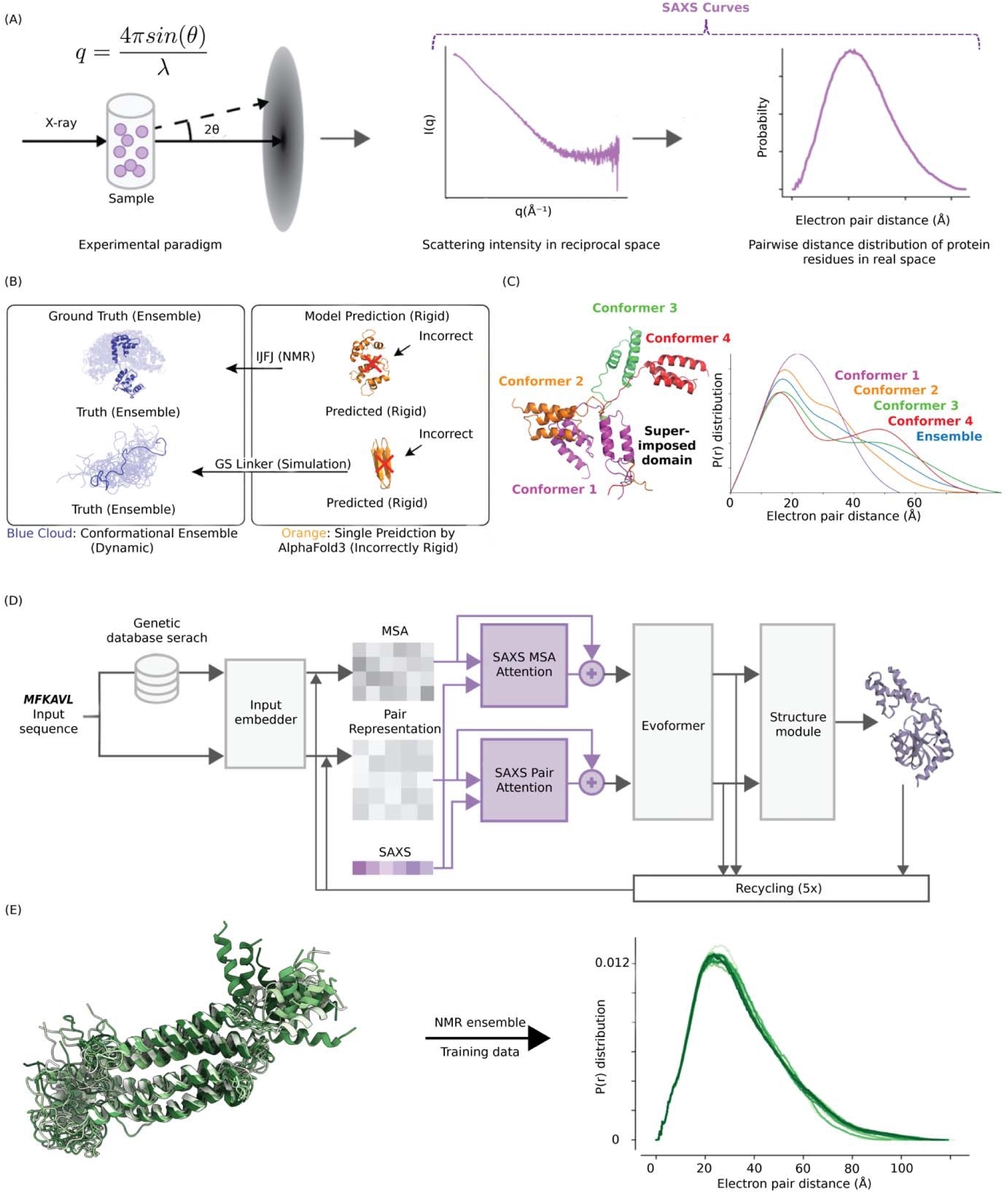
Integrating SAXS into Protein structure predictions. **(A)** SAXS captures structural details of proteins as they exist in a physiologically-relevant solution environment. (Left: The Diagram of SAXS experiment. Middle: Reciprocal space data of SAXS. Right: Pair-wise distance distribution P(r) derived from SAXS.) **(B)** AlphaFold3 demonstrates reduced accuracy in modeling ensembles of conformations, as found for proteins with highly flexible or disordered protein regions under solution environments. (Top: NMR ensemble of 1JFJ and AlphaFold3 prediction, Bottom: GS16 ensemble simulation^19^ and AlphaFold3 prediction) **(C)** SAXS pair-wise distance information can reveal the difference between different protein conformations. **(D)** The AlphaSAXS architecture utilizes SAXS as a native input for protein structure prediction by integrating experimental SAXS profiles with the latent MSA and pair representations. Model weights were fully retrained to convergence on a customized dataset, using the original AlphaFold2 weights as an initial baseline. **(E)** AlphaSAXS is trained with solution NMR ensembles and synthesized SAXS profiles that recapture the solution protein conformations under biological environment.

Here, we present AlphaSAXS, a novel approach for integrating experimental SAXS data with protein structure prediction with a customized curated dataset. AlphaSAXS augments the OpenFold architecture,^18^ incorporating SAXS pairwise distance distributions as additional model input to guide structural inference. By leveraging physiologically relevant experimental structural information, AlphaSAXS demonstrates improved accuracy and conformational flexibility over sequence-only models. We evaluate its performance on apo-holo protein pairs and newly collected NMR ensembles, demonstrating that our method guided by SAXS successfully generates distinct conformations for the same sequence that match experimental scattering curves with high fidelity. AlphaSAXS thus provides a foundation for a fully automated, high-throughput SAXS modeling pipeline.

## Results

### Small-Angle X-ray Scattering as a Structural Constraint

SAXS is a high-throughput, solution-based technique ideally suited for the rapid assessment of global protein topology and conformational ensembles (Fig. 1A). Previous studies, notably by Baker and colleagues,^20^ have demonstrated the utility of SAXS in validating and refining computationally designed proteins. Because the structural discrepancies manifest as significant deviations in the pair distance distribution function, *P*(*r*), SAXS profiles serve as a sensitive diagnostic for inaccuracies in predicted model conformations. For instance, structural hypotheses generated via diffusion-based sampling in the latest AlphaFold3 architecture fail to recapitulate the dynamic ensemble of the calcium-binding protein 1JFJ.^21^ Furthermore, AlphaFold3 exhibits reduced predictive accuracy on intrinsically disordered regions, such as the GS16 protein linker (Fig. 1B). Conversely, SAXS *P*(*r*) profiles can robustly distinguish between distinct conformational states of a single sequence, as demonstrated by the critical assessment of protein structure prediction (CASP) target T1200, which serves as an experimental benchmark for AI model evaluation (Fig. 1C).^22^ Finally, similar to the latent pair representation, which encodes distogram probability matrices, the reciprocal SAXS profile captures information about the pairwise distances between atoms. However, SAXS resolves distances much more finely and has no maximum distance limit. Thus, by integrating these experimental constraints directly into the AlphaFold/OpenFold architecture,^18^ our proposed AlphaSAXS model addresses the inherent limitations of current fold-prediction algorithms, providing the critical information necessary to correct erroneous structural hypotheses.

### Architectural Integration of SAXS profiles

Successfully guiding existing AI-driven models with experimental SAXS data requires innovation of both the model architecture as well as advancement of the training dataset and training strategy to incorporate SAXS data and multiple conformations for the same sequence.

First, we developed a modified architecture capable of processing different SAXS curve inputs from different conformations of a single protein sequence, thereby enabling the prediction of distinct conformational states with the same sequence but different SAXS profiles (Fig. 1D).

More specifically, we introduced SAXS-derived information into two of the network’s primary internal latent representations: the pair representation and the MSA representation. The modification of MSA representation has been proven to alter the model prediction and thus a SAXS conditioned MSA network is a reasonable approach for reconstructing the structure based on SAXS.^23^ In addition, while the pair representation encodes the spatial relationships between residue pairs, the SAXS scattering intensity provides an experimental proxy for the global electron distance distribution with the *P*(*r*). By aligning these features, AlphaSAXS can refine the initial spatial predictions based on experimental distance constraints and thus we can leverage this newly modified architecture to allow the model to explore the conformational landscape guided by SAXS profiles.

The above design is achieved through two multi-headed cross-attention modules: **SAXS-MSA-Attention** and **SAXS-Pair-Attention**. In these modules, the MSA and pair embeddings serve as queries to the SAXS profile. The resulting attention weights determine the influence of specific SAXS intensity bins on individual MSA clusters and residue pairs. This mechanism allows the model to dynamically update its internal embeddings based on the most relevant features of the experimental profile. While the modulation of MSA attention has been previously demonstrated to influence AlphaFold/OpenFold predictions^23^, the SAXS-Pair-Attention module represents a novel contribution, explicitly leveraging the rich pairwise distance information encoded within the SAXS profile.

By integrating experimental evidence with the common latent representations of the AlphaFold/OpenFold model, this approach allows us to enhance and leverage the strength and proven architecture of AlphaFold, rather than replace it. This strategy further has the advantage that it provides a reusable design, enabling the integration of our SAXS-MSA and SAXS-Pair modules with other structure prediction models that utilize similar latent representations as well the integration of additional experimental evidence and modalities with our architecture in a similar fashion in the future.

### Ensemble-Based Training Strategy

The development of AlphaSAXS required the creation of a specialized training dataset that supports the prediction of diverse, multi-state structures (Fig. 1E). A significant limitation of AlphaFold/OpenFold arises from its training on X-ray crystallography, cryo-EM structures, where flexible regions are often unresolved or omitted in the PDB. While AlphaFold2 incorporates Nuclear Magnetic Resonance (NMR) structures, it typically utilizes only the first model of an ensemble^1^, enforcing a rigid “one sequence, one structure” paradigm.

To overcome this limitation, we retrained our model using a curated collection of over 10,000 NMR ensembles from the PDB, each providing 20–30 diverse conformations per protein, comprising over 200,000 individual training samples in total. This shifts the training objective toward a “one sequence, multiple conformations” relationship. To prevent data leakage, we rigorously excluded any proteins sharing greater than 70% sequence similarity with our test dataset during the training process.

Furthermore, we developed custom functions to generate synthetic SAXS data for each conformation. By pairing these unique SAXS profiles with their corresponding structural members within an ensemble, AlphaSAXS learns to generate distinct conformations from the same primary sequence. Consequently, the model can predict structural transitions driven by biological or environmental conditions—as captured by SAXS—rather than relying solely on sequence encoding and evolutionary information.

### Benchmarking AlphaSAXS on Conformational Transition Targets

AlphaSAXS was specifically designed to resolve structural targets that lie beyond the predictive capacity of the standard OpenFold architecture. To evaluate its performance, we curated a test set of 40 proteins from the Apo-Holo pair dataset, representing documented failure modes in standard AlphaFold/OpenFold predictions^24^. Following a training regime of over 3,000 GPU hours, we benchmarked the model’s accuracy against the baseline OpenFold architecture.

Our evaluation revealed that AlphaSAXS achieves superior performance, demonstrating an overall Root-Mean-Square Deviation (RMSD) improvement of 0.08 Å across the dataset, with improvements exceeding 2.0 Å for targets that OpenFold failed to predict accurately. Additionally, the SAXS *L*1 loss improved by more than 20% on average, and by over 50% on these challenging targets (Fig. 2A, B, C). We also observed a similar improvement on the radius of gyration (*R_g_*) of predicted proteins (Fig. 2A). This is based on categorizing the test set into three performance tiers depending on the OpenFold baseline RMSD: high-accuracy (≤ 1.0 Å), mid-accuracy (1.0 – 5.0 Å), and low-accuracy (> 5.0 Å). AlphaSAXS demonstrated the most significant accuracy gains on the low-accuracy targets where OpenFold previously failed, while maintaining comparable performance—with only marginal fluctuations—on targets that were already accurately predicted by the baseline (Fig. 2B, C).

**Figure 2.**
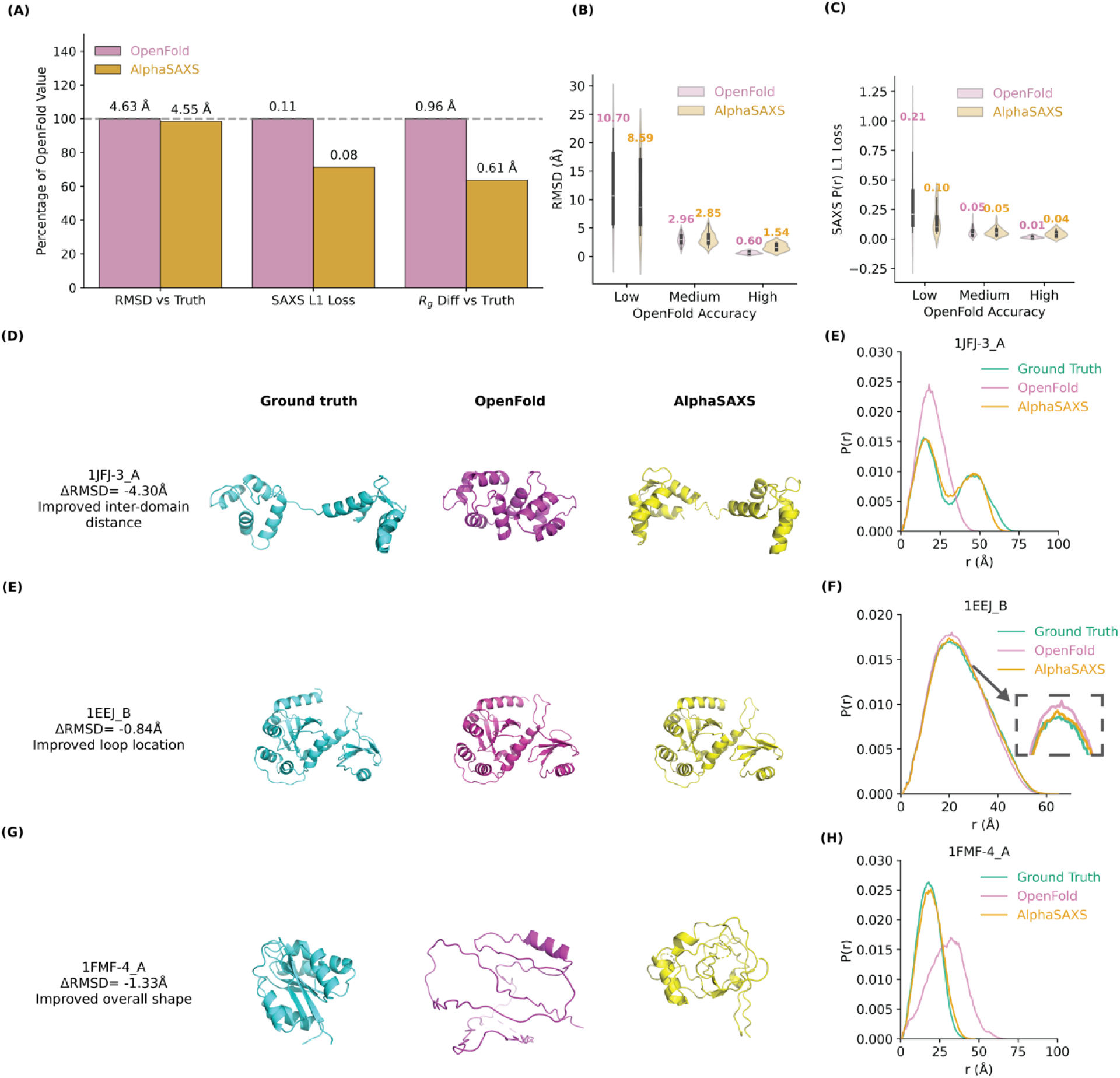
AlphaSAXS improves the conformational prediction accuracy of OpenFold. **(A)** AlphaSAXS outperforms the OpenFold baseline, even when restricted to a smaller training dataset, achieving an overall 0.03 reduction in SAXS L1 loss, a 0.08 Å improvement in RMSD and a 0.35Å improvement in the radius of gyration (R_g_). **(B, C)** AlphaSAXS maintains high stability, preserving accuracy for structures already well-predicted by OpenFold while significantly improving performance on challenging targets where the baseline fails. OpenFold baseline accuracy tiers (high, mid, low) are defined by prediction RMSD thresholds of 1.0 Å and 5.0 Å. **(D, E)** AlphaSAXS successfully recaptures the elusive expanded conformation of the calcium-binding protein 1JFJ, accurately resolving the binding pocket. **(F, G)** Improved loop prediction accuracy for the protein disulfide-bond isomerase 1EEJ. The correct loop conformations recovered by AlphaSAXS are crucial for biological dimerization. **(H, I)** AlphaSAXS improves the prediction of the overall structural topology for 1FMF, a glutamate mutase from *Clostridium tetanomorphum*. Due to its origin from a rare species, the MSA for 1FMF exhibits weak coverage, leading to an incomplete fold prediction by OpenFold. By integrating SAXS data, AlphaSAXS successfully guides the model to form the correct overall shape and size.

To elucidate the biophysical implications of these improvements, we examined three representative case studies: the calcium-binding protein 1JFJ,^21^, the isomerase 1EEJ^25^ and 1FMF,^26,27^ a rare protein from *Clostridium tetanomorphum*.

For **1JFJ**, standard AlphaFold and OpenFold predictions are poor, typically generating an overly compact structure that deviates significantly from NMR-derived ensembles (Fig. 2D, E). By providing a reference SAXS profile as a structural guide, AlphaSAXS successfully recaptures the expanded conformational state observed in the reference PDB. This capability underscores the potential for AlphaSAXS to uncover physiologically relevant poses that are inaccessible to sequence-only models.

In the case of **1EEJ**, while OpenFold accurately predicts the active site, it fails to resolve the flexible loop, which is critical for dimerization and essential for its biological function (Fig. 2F, G). AlphaSAXS achieved a precise refinement of this specific loop, improving the overall RMSD by 0.84 Å. This result suggests that AlphaSAXS can extract not only global topological constraints from SAXS profiles but also fine-grained structural details that govern protein-protein interactions.

The protein **1FMF** presents a unique challenge due to low evolutionary coverage (poor MSA depth) (Fig. 2H, I). Standard OpenFold fails to generate a folded conformation for this protein. In contrast, AlphaSAXS generated a partially folded structure with a *R_g_* consistent with experimental data. This case study highlights a critical nuance: while SAXS profiles can provide essential global geometric constraints to stabilize folding, they cannot entirely compensate for deficits in MSA information on rare species and proteins within the inherent OpenFold mode and training dataset.

Taken together, these examples demonstrate that the observed improvements in RMSD translate to substantial gains in predicting biologically relevant structural features. Importantly, these corrections are achieved not through rigid-body post-processing, but by dynamically modulating the network’s internal representations during the folding trajectory. These findings confirm that low-resolution SAXS data can be used to redirect the AlphaFold/OpenFold architecture’s structural inference, successfully forcing its predictions to escape local minima and align with the experimental solution-state conformation. They also validate that our SAXS-driven architecture successfully learned loss functions as intended, effectively bridging the gap between sequence-based prediction and solution-state experimental reality.

### Generating Distinct Conformations from an Invariant Sequence

In biological systems, a single protein sequence can adopt disparate conformations depending on ligand binding, complex formation, or environmental fluctuations—changes that are often reflected in the SAXS profile. AlphaSAXS is uniquely positioned to address the “one sequence, one structure” limitation of traditional models by generating distinct, experimentally grounded conformations. This approach not only provides a solution to the bias of OpenFold towards native PDB structures but also ensures that the generated states remain within the realm of physically plausible structures, avoiding the out-of-distribution artifacts sometimes observed in sequence-only generative models.^28^

To demonstrate this concept, we utilized the 1JFJ/1JFK apo-holo pair which shared the same sequence but different conformations (Fig. 3A). We synthesized two distinct *P*(*r*) curves corresponding to the ground-truth apo and holo states. Guided by these disparate SAXS curves, AlphaSAXS generated two distinct structural models with an inter-state RMSD > 4 Å, effectively capturing the large-scale conformational shift. Conversely, for the 1EEJ sequence (Fig. 3B), the apo and holo states yield nearly identical *P*(*r*) profiles. In this instance, AlphaSAXS predicted highly similar structures for both inputs, accurately reflecting the lack of global structural divergence indicated by the experimental data.

**Figure 3.**
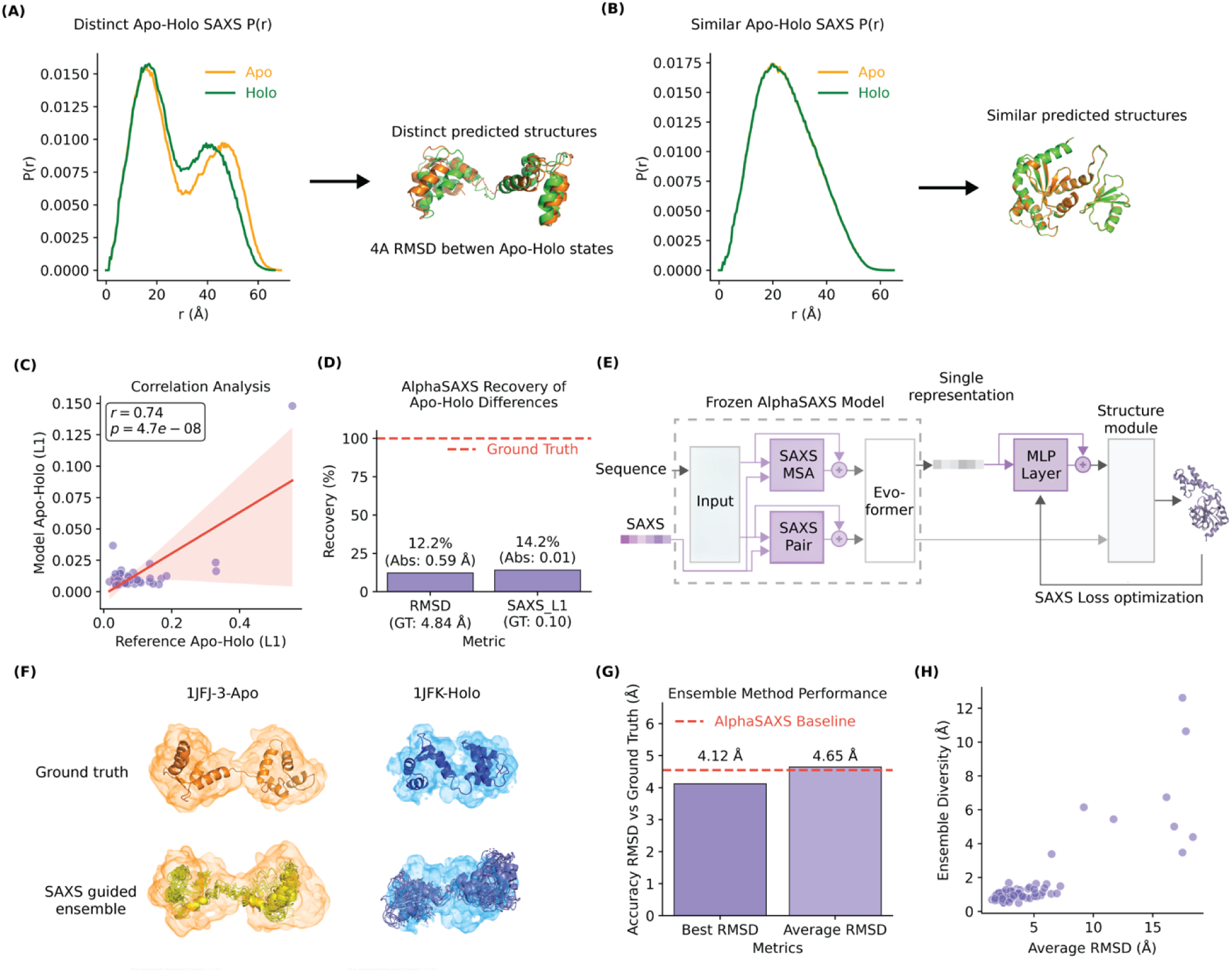
Evaluating the conformational diversity and ensemble performance of AlphaSAXS. **(A)** AlphaSAXS generates distinct conformational states for the apo-holo pair 1JFJ/1JFK based on disparate SAXS profiles. **(B)** AlphaSAXS yields indistinguishable structural ensembles when provided with highly similar P(r) profiles (e.g., 1EEJ). **(C)** Correlation between input P(r) divergence and the resulting structural diversity of output models. **(D)** Recovery of experimental structural variance; AlphaSAXS recovers approximately 15% of the conformational difference between the apo-holo state with the same sequence input. **(E)** Architectural schematic of the inference-time optimization protocol for ensemble generation. **(F)** Representative apo-holo structural ensemble. Shaded volumes represent the estimated electron density derived from the synthesized SAXS P(r) profile. **(G)** RMSD accuracy of generated ensembles relative to ground-truth structures for the test dataset. AlphaSAXS ensemble has a 0.53Å RMSD improvement over the OpenFold predictions. **(H)** Correlation between intra-ensemble conformational diversity and structural accuracy across target groups.

To quantify this behavior, we analyzed the correlation between the *L*1 difference of input *P*(*r*) pairs and the resulting output diversity of AlphaSAXS. We observed a positive correlation (Pearson’s *r* = 0.57), indicating that as the experimental evidence for structural change increases, AlphaSAXS accordingly scales its conformational output (Fig. 3C). However, we note that the generated structural diversity typically captured less than 20% of the theoretical ground-truth difference (Fig. 3D). Thus, this test reveals that, while the AlphaSAXS model is responsive to SAXS profiles when conformational changes are large, to fully sample the expansive conformational states of highly dynamic proteins requires further modification and better sampling strategy on the model.

### Enhancing Diverse Conformation Prediction via Inference-Time Optimization

We hypothesized that the difficulty in exploring the full conformational landscape of a single protein stems from high-energy barriers between dynamic states. To address this, we developed an inference-time optimization protocol that utilizes a SAXS-specific loss to steer AlphaSAXS predictions toward the experimental target. This is achieved by implementing a lightweight, multi-layer adaptation network situated between the Evoformer layers and the final structure module, which generates small perturbations guided by the SAXS loss during optimization (Fig. 3E). To prevent disruption of the learned representations, all primary AlphaSAXS weights remain frozen during inference-time optimization, while the weights of the adaptation network are fine-tuned to align the structural output with the input *P*(*r*) curve.

We evaluated this strategy using the test dataset described above. By ranking candidate structures based on their SAXS loss, we predicted ensembles consisting of 50 conformations for each of the 40 proteins in the Apo-Holo dataset. Evaluation of these ensembles revealed that the best-fitting conformation achieved an RMSD improvement of approximately 0.4 Å over the single-state model (Fig. 3F). Moreover, the average RMSD of the ensemble members remained on par with the baseline model, suggesting that the adaptation network generates diverse conformations without sacrificing local structural quality.

To quantify the diversity of these conformations, we analyzed the intra-ensemble RMSD. We observed significant internal heterogeneity, with most ensembles exhibiting an internal RMSD exceeding 1.0 Å, and some surpassing 4.0 Å (Fig. 3H). This diversity indicates that the adaptation network does not merely produce redundant samples but actively explores the conformational space to satisfy the experimental *P*(*r*) profile. By integrating this protocol, we have established a robust candidate-generation framework capable of translating a primary sequence and a SAXS profile into a biologically relevant structural ensemble.

### Integrating AlphaSAXS Output with Hydration Layer Modeling

Experimental SAXS profiles inherently reflect contributions from the protein’s hydration shell (water layer), a component typically addressed during post-processing using algorithms such as FoXS, CRYSOL, WAXSIS, or AQUASAXS.^29–32^ As AlphaSAXS was trained on synthetic data without explicit water modeling, it is essential to validate the compatibility of its generated ensembles with these standard biophysical workflows. To evaluate this, we utilized the Staphylococcal protein A target (T1200) from the CASP16 ensemble prediction category (Fig. 4A). It is important to note that while CASP competitors address the significantly harder challenge of blind prediction without experimental scattering data, our objective here is distinct: we utilize the experimental SAXS profile specifically to validate AlphaSAXS as a viable sampling method for hydration layer modeling.

**Figure 4.**
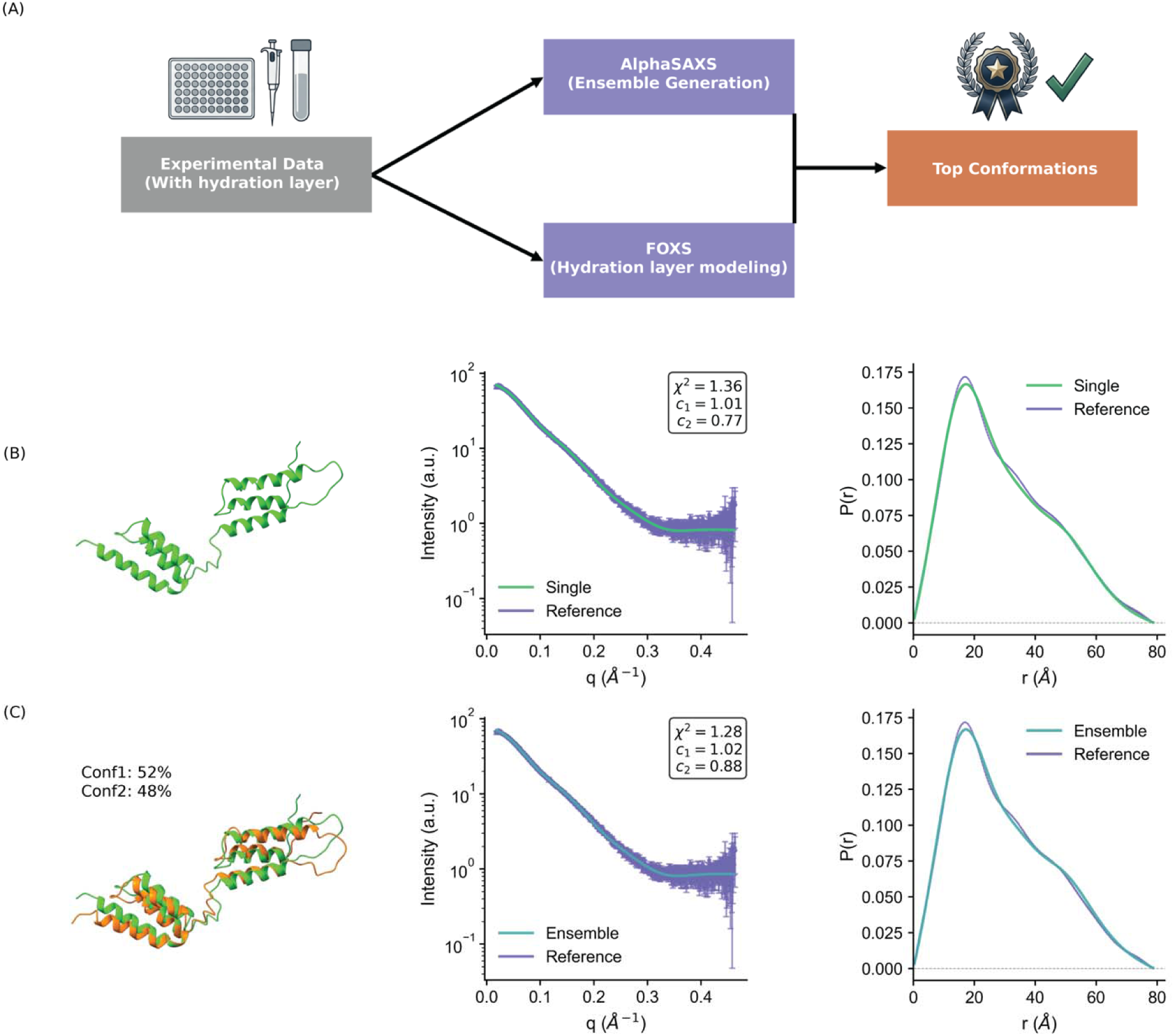
Modeling the protein hydration layer with AlphaSAXS ensembles. **(A)** Schematic protocol for modeling AlphaSAXS ensembles using the FoXS algorithm based on experimental data. **(B)** Accurate hydration layer modeling for the T1200 protein, utilizing FoXS to select the optimal conformation from the AlphaSAXS-generated ensemble. **(C)** Hydration layer modeling with MultiFoXS, demonstrating the selection of a weighted ensemble of conformations to fit the experimental profile.

We employed the experimental SAXS profiles to guide AlphaSAXS to generate candidate ensembles for the Protein A wild-type target T1200, subsequently using FoXS to model the hydration layer and identify the optimal conformation. The best single model selected by FoXS yielded a *χ*^2^ value of approximately 1.3, indicating a high-accuracy prediction consistent with experimental data.^29,33^ Furthermore, we employed MultiFoXS to select a weighted minimal ensemble, which also demonstrated excellent agreement with the experimental profile (Fig. 4B, C).^33^ These results confirm that coupling AlphaSAXS ensemble generation with FoXS hydration modeling achieves accuracy on par with established protocols such as BilboMD. However, we caution that the generated ensemble may not fully represent the true thermodynamic ensemble of Protein A due to the absence of NMR-based training and verification. Consequently, we position our method as an enhanced sampling technique, analogous to Molecular Dynamics (MD) simulations, rather than a definitive ensemble generator for blind predictions.

## Discussion

### Importance of conformational change

The work in this paper is an important theoretical advance in linking AI structure prediction to non-human interpreted experimental data. While AI structure prediction algorithms have achieved remarkable levels of accuracy compared to a decade ago^34^ significant hurdles remain for reliable mechanistic interpretation and small molecule discovery. A major constraint is the difficulty in predicting conformational heterogeneity especially in biological samples in solution conditions.^35^ This is a critical limitation because conformational changes, ranging from active site loops to large interdomain movements, are the primary mechanism regulating protein function in the biological environment. Many current algorithms are trained to ignore these physiological fluctuations, effectively associating a single sequence with static structures. In contrast, SAXS profiles inherently capture structural responses to the solution environment. By leveraging these experimental constraints, this work demonstrates that it is indeed possible to retrain the underlying architecture to successfully handle the “one sequence, multiple structures” paradigm.

### The importance of the experimental driven prediction

There are certain recently developed algorithms targeted at solving the previously mentioned one-sequence multiple structure problem like BioEmu.^36^ However, these algorithms are primarily trained on MD simulations which have a high probability of not matching the experimental data during the last CASP in the ensembles category.^22^ Thus here we discussed the importance of incorporating experimental data directly into the training and input process of the model.

One step towards increasing knowledge of dynamicity is the inclusion of experimental data, not as a filter, but as part of the assessment cycle. Use of experimental data to identify AI structure predictions consistent with that data has been tested on crystals, with success.^37,38^ However, for the AI model to incorporate the structural information, the algorithm must learn that its predictions are “wrong” and are not consistent with the experimental data. Here, we show that we can employ SAXS information as part of the training loss function and alter the output predictions.

It must be carefully differentiated that AI structure prediction algorithms use atomic models based on experimental data, but do not use the uninterpreted experimental data that reflects the dynamic range of the molecule, from large conformational changes to atomic level movements. Recently, algorithms altering the MSA or inclusion of molecular dynamics models have attempted to bypass these limitations. MD is powerful, but is purposefully biased towards compaction to prevent the fold from exploding. Thus, inclusion of our SAXS loss function, reflecting the true solution structure, was able to push out the structure prediction from local minima.

### Future Directions and Limitations

While AlphaSAXS establishes a robust framework for experimental guidance, future developments must address the complexities of the solvent environment. Currently, our model does not explicitly account for the hydration layer during training, relying instead on post-processing tools like FoXS. In the future, we need to include a hydration layer to make it more accurately useful against experimental SAXS data. The hydration layer for disordered regions will be linear with mass of the disordered region, but is not linear with folded regions. In theory, future development of AlphaSAXS could take advantage of this hydration layer nuance. Since the hydration shell scales linearly with mass for disordered regions but non-linearly for folded domains, future iterations of AlphaSAXS could exploit this nuance to better differentiate between ordered and disordered states directly from the scattering profile. Furthermore, the principles established here, guiding inference with global and uninterpreted constraints are broadly applicable. Extending this approach to include raw data from crystallographic or CryoEM densities or from NMR chemical shifts promises to further refine our ability to predict protein dynamics, ultimately enabling the reliable use of structure prediction models for predictive biology.

## Methods

### SAXS-guided Attention Modules

To integrate SAXS profiles while leveraging the existing AlphaFold architecture, we introduced SAXS experimental data into the two key inputs to the network, the pair representation and MSA. We targeted the pair representation because SAXS profiles reflect pairwise distances between residues, and similarly, the pair representation encodes information about the relationship between pairs of residues. We also targeted the MSA representation because experimental SAXS profiles often contain information from a mixture of multiple conformations in solution, and previous work has demonstrated that modifying the MSA during inference can be used to sample alternative conformational states. To incorporate SAXS information with these inputs, we added two multi-headed cross-attention modules, the SAXS-MSA-Attention and SAXS-Pair-Attention modules, to perform attention between the MSA and pair input embeddings and the p(r) probability distribution from the SAXS profiles. Specifically, the pair and MSA representations are used to form the queries for the SAXS profile, and the attention weights are calculated to determine how each SAXS bin might influence each MSA cluster and residue or residue pair, allowing these input embeddings to be updated based on relevant SAXS profile features. In the SAXS-MSA-Attention block, the number of heads was *N_heads_* = 8, and the dimension of the keys, queries, and values was *c* = 32, and in the SAXS-Pair-Attention block the dimensions were *N_heads_* = 4, *c* = 32. The output of these modules was added to the MSA and pair representational inputs to the Evoformer.

### Training Data

We trained this network using training data with structures with multiple conformations. We fine-tuned the weights of our AlphaSAXS model starting with the publicly available pre-trained OpenFold for the architecture implementation, weight, and training pipeline. For our NMR dataset, we obtained over 12,000 samples of NMR models from the PDB. We removed any sequences with more than 70% similarity to the test dataset. We separated the conformations for NMR data with the BioPython library to obtain a training dataset. For initial training, we filtered out recordings with a number of residues > 256 since AlphaFold used a similar constraint^1^ and since during training the residue dimension is randomly cropped to length *N_res_* = 256, and we wanted to ensure our SAXS distributions corresponded to the cropped structures without recalculating during training. We used ColabFold search with the MMSeqs2 clustering suite to generate the MSAs used in training with our in-house server.

### Target Dataset

For model evaluation, we used a subset of the curated collection of apo-holo pairs of protein conformers by Saldaño et al., 2022, which provides two unique conformations for the same protein under different physiological processes. The initial collection of 91 pairs was filtered to exclude sequences with a *N_res_* > 256. To focus our analysis on proteins that did not already have highly accurate predictions by the original AlphaFold, we selected protein pairs with a total pair *RMSD* > 4Å and an individual conformer *RMSD* > 2.5Å, resulting in an evaluation dataset of 40 proteins. We calculated the average Cα-RMSD change of the conformation unfavoured by AlphaFold for all protein sequences in the target dataset. Compared to the OpenFold, our model generally shows an improvement in average RMSD and lower standard deviation

### SAXS profile generation

The experimental SAXS profile is limited with the possible water layer problem and non-standard format. For effective training of our model on a larger training dataset, we decided to generate synthetized SAXS dataset based on existing algorithms. We calculated the distance between each atom of the given PDB and then converted it into a pairwise distance distribution of electrons as provided in SAXS data. The algorithm has been tested against widely used SAXS software, including FOXS, GNOM, and RAW. The SAXS profiles for both the training dataset and the test dataset are prepared ahead of time using GNU parallel for each PDB. The SAXS profile of the predicted structure for the loss function calculator or inference optimization is calculated on the fly using an embedded version of the same algorithm.

### AlphaSAXS Loss Function

To train AlphaSAXS, we augmented the standard OpenFold loss function with a differentiable SAXS *L*_l_ loss term. The composite loss function is defined as:

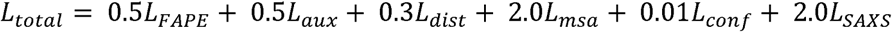

where *L_FAPE_*, *L_aux_*, *L_dist_*, *L_msa_*, and *L_conf_* correspond to the loss components defined in the original AlphaFold2 architecture, utilizing identical weighting. The SAXS term, *L_SAXS_*, represents the *L*_l_ distance between the predicted and experimental *P*(*r*), utilizing a custom differentiable implementation weighted to balance the magnitude of the structural loss components within the overall optimization landscape.

### Inference-Time Optimization

To generate diverse structural ensembles, we implemented an inference-time optimization protocol that modifies the AlphaSAXS architecture by inserting a two-layer Multilayer Perceptron (MLP) adapter between the EvoFormer stack and the final Structure Module. During this optimization phase, all pre-trained AlphaSAXS weights are frozen. The MLP adapter weights are iteratively updated via gradient descent, minimizing the SAXS loss of the predicted structure relative to the target profile. The optimization proceeds for a maximum of 500 iterations, with early stopping triggered if the SAXS loss fails to improve for 50 consecutive iterations. The trajectory is recorded, and the top 50 structures exhibiting the lowest SAXS loss are retained as the final ensemble for metric evaluation.

### Evaluation and Visualization

Structural comparison between predicted models and ground-truth references was performed using custom scripts based on the MDTraj analysis suite. Source code for evaluation metrics and visualization pipelines is available in the associated GitHub repository.

### Hydration Layer Modeling

For validation against experimental scattering data, we employed FoXS to explicitly model the hydration shell for each conformation within the AlphaSAXS-generated ensembles. Briefly, FoXS calculates the theoretical scattering profile by adjusting the contrast of the hydration layer and the excluded volume via two fit parameters, *c*_l_ and *c*_2_. Model quality was assessed using the discrepancy (*χ*^2^) between the calculated and experimental profiles. The conformation yielding the minimal *χ*^2^ value was selected for structural analysis and visualization

## Supporting information

PDB structure and SAXS curve generated by AlphaSAXS

## Data Transparency

The model code and weight is stored in the Github repo https://github.com/lbl-cbg/metfish. The training dataset and test dataset will be available upon request. The PDB structure and SAXS curve generated by AlphaSAXS is attached to the supplemental materials.

## Acknowledgements

We thank Frank DiMaio and John Tainer for their helpful discussions on our training databases and AI model. This work was supported by the Laboratory Directed Research and Development Program of Lawrence Berkeley National Laboratory under U.S. Department of Energy Contract No. DE-AC02-05CH11231 (S.T. and O.R.), NIH R01 GM137021 (S.T.), and NIH R35GM150982 (K.P.). This research used resources of the National Energy Research Scientific Computing Center (NERSC), a Department of Energy User Facility using NERSC award AI4Sci@NERSC-ERCAP0034111.

