## Supplementary material for "Experimental Data Driven AI Framework for Flexible Protein Conformational Reconstruction": PDB structure and SAXS curve generated by AlphaSAXS

### Supplementary Figure - Page 1/20

Ground Truth

OpenFold

AlphaSAXS

P(r)

--- 1AEL-12\_A (Apo) 1URE-8\_A (Holo) ---
OpenFold
RMSD=2.49 Å AlphaSAXS
RMSD=2.68 Å

Ground Truth

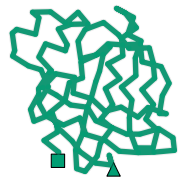

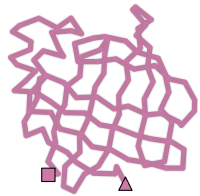

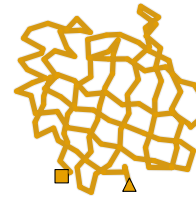

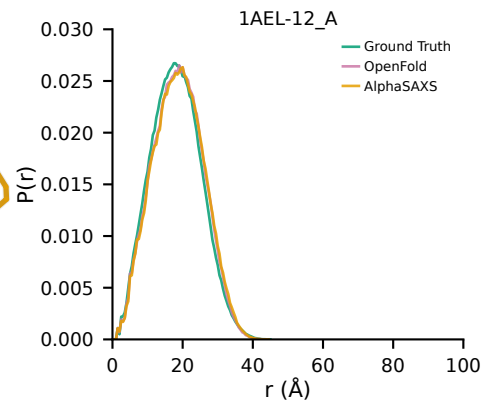

Ground Truth

OpenFold
RMSD=1.50 Å

AlphaSAXS
RMSD=1.61 Å

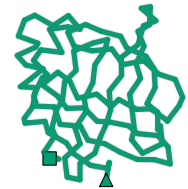

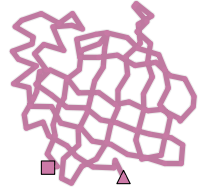

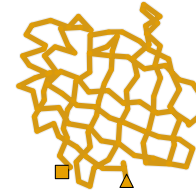

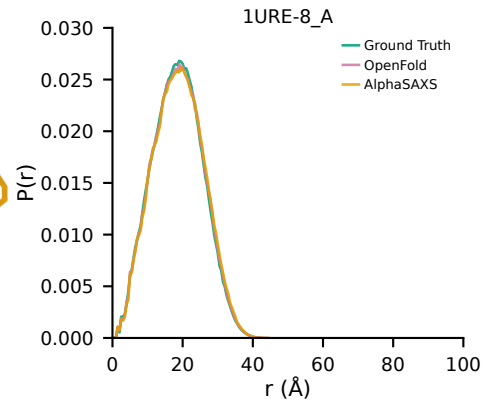

Ground Truth

--- 1EX6\_B (Apo) 1EX7\_A (Holo) ---
OpenFold
RMSD=1.32 Å AlphaSAXS
RMSD=3.10 Å

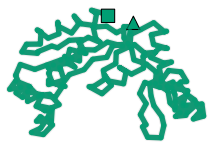

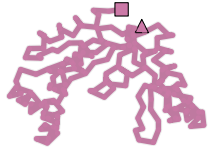

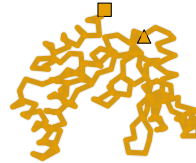

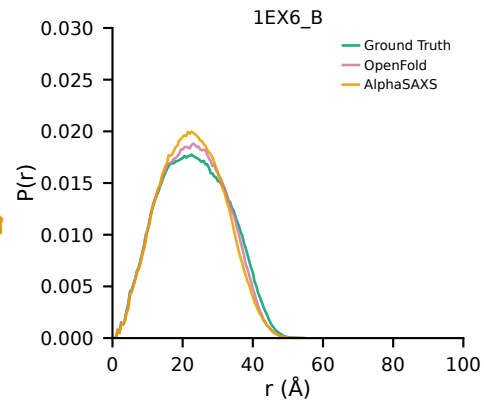

Ground Truth

OpenFold
RMSD=3.40 Å

AlphaSAXS
RMSD=2.17 Å

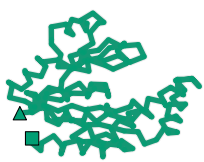

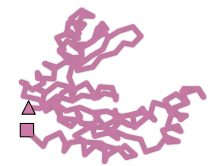

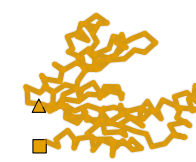

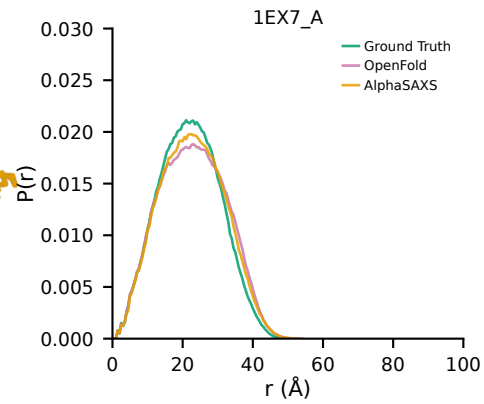

### Supplementary Figure - Page 2/20

Ground Truth

OpenFold

AlphaSAXS

P(r)

--- 1F3Y-17\_A (Apo) 1JKN-15\_A (Holo) ---

OpenFold
RMSD=3.32 Å

AlphaSAXS
RMSD=5.41 Å

Ground Truth

Apo

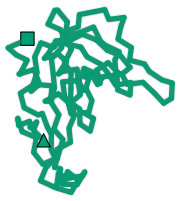

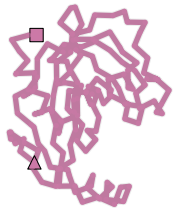

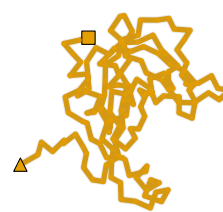

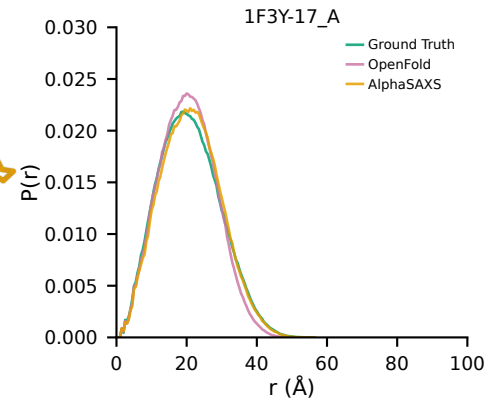

Ground Truth

OpenFold
RMSD=2.73 Å

AlphaSAXS
RMSD=4.15 Å

Holo

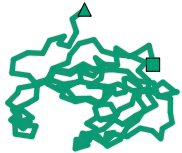

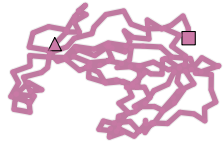

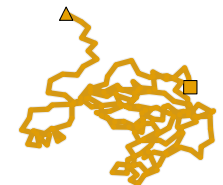

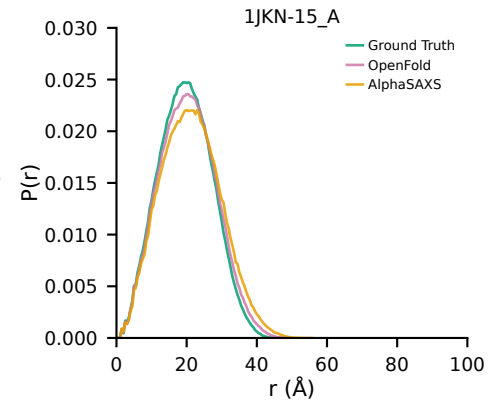

Ground Truth

--- 1FMF-4\_A (Apo) 1ID8-11\_A (Holo) ---

OpenFold
RMSD=17.03 Å

AlphaSAXS
RMSD=15.70 Å

Apo

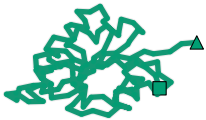

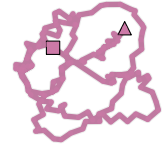

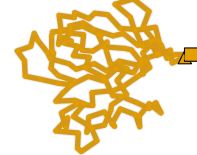

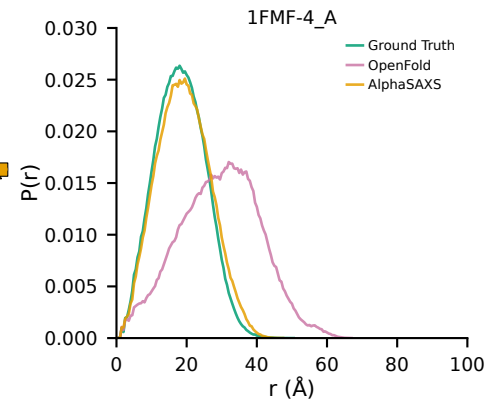

Ground Truth

OpenFold
RMSD=17.52 Å

AlphaSAXS
RMSD=16.62 Å

Holo

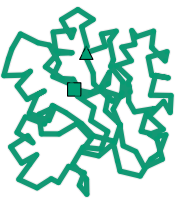

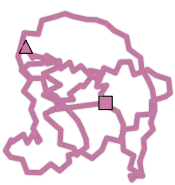

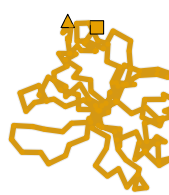

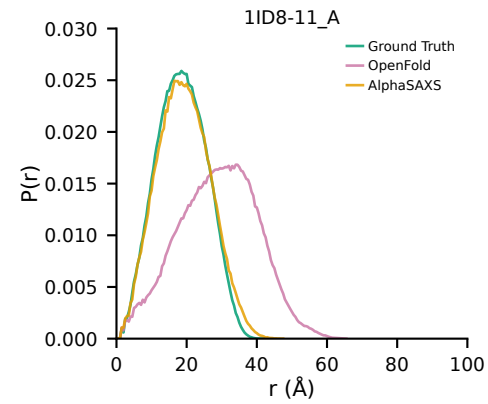

### Supplementary Figure - Page 3/20

Ground Truth

OpenFold

AlphaSAXS

P(r)

--- 1GH1-7\_A (Apo) 1CZ2-8\_A (Holo) ---

OpenFold
RMSD=2.54 Å

AlphaSAXS
RMSD=2.32 Å

Ground Truth

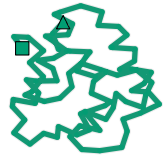

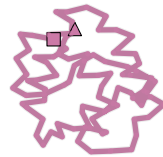

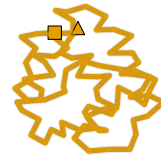

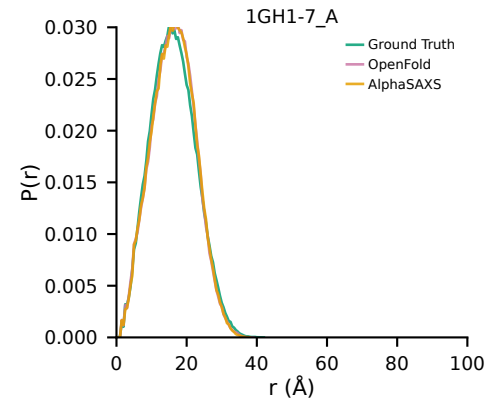

Ground Truth

OpenFold
RMSD=2.69 Å

AlphaSAXS
RMSD=2.43 Å

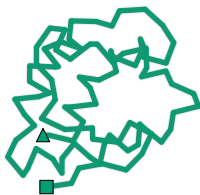

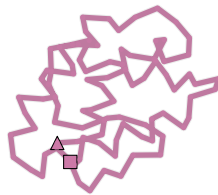

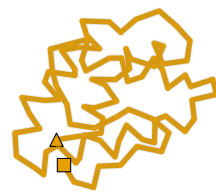

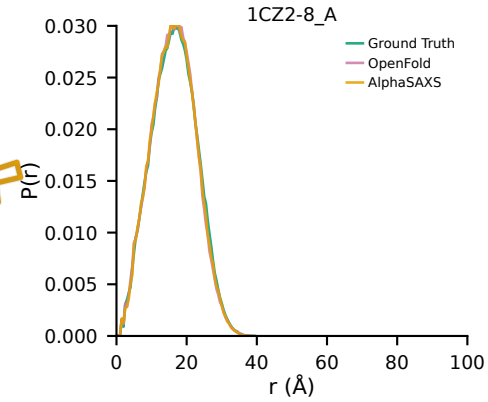

Ground Truth

--- 1IJA-15\_A (Apo) 2KID-4\_A (Holo) ---
OpenFold
RMSD=3.42 Å

AlphaSAXS
RMSD=4.38 Å

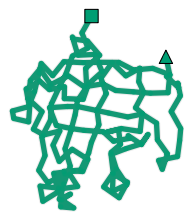

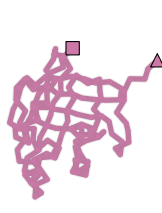

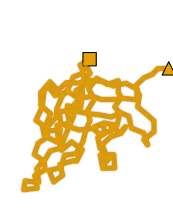

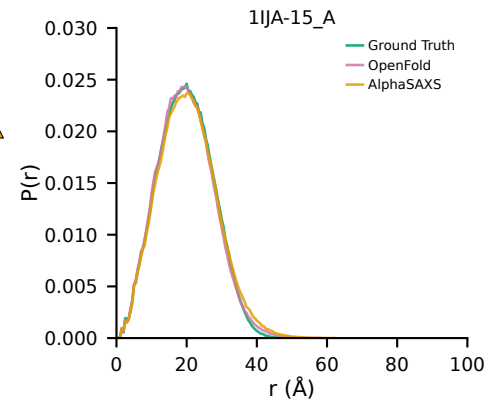

Ground Truth

OpenFold
RMSD=2.34 Å

AlphaSAXS
RMSD=1.89 Å

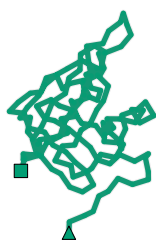

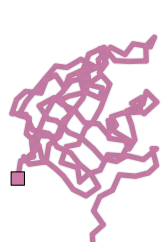

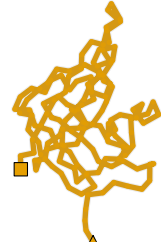

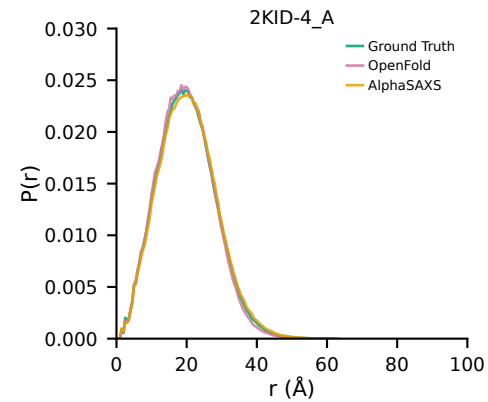

### Supplementary Figure - Page 4/20

Ground Truth

OpenFold

AlphaSAXS

P(r)

--- 1JFJ-3\_A (Apo) 1JFK\_A (Holo) ---

OpenFold
RMSD=13.63 Å

AlphaSAXS
RMSD=9.33 Å

Ground Truth

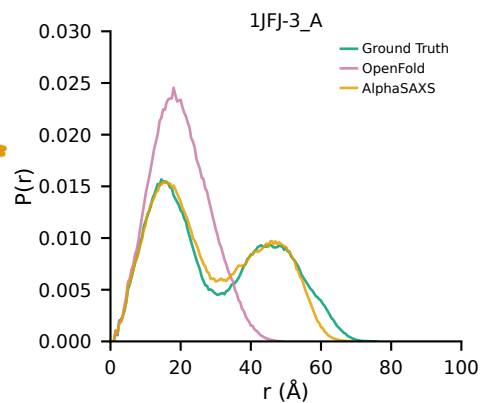

Ground Truth

OpenFold
RMSD=13.27 Å

AlphaSAXS
RMSD=10.84 Å

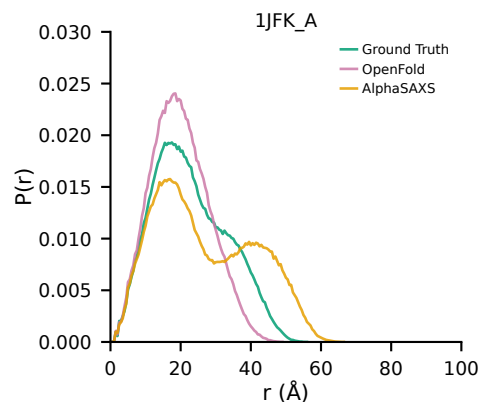

Ground Truth

--- 1JM4-15\_B (Apo) 1WUM\_A (Holo) ---
OpenFold
RMSD=2.97 Å

AlphaSAXS
RMSD=4.24 Å

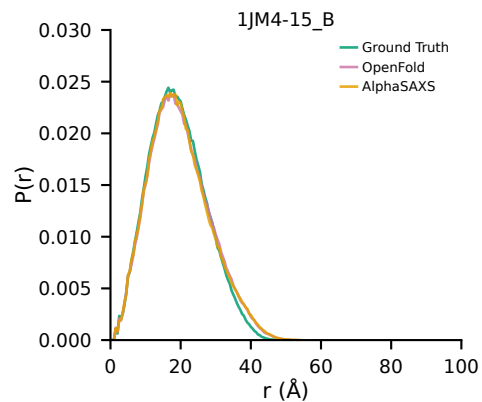

Ground Truth

OpenFold
RMSD=4.02 Å

AlphaSAXS
RMSD=5.19 Å

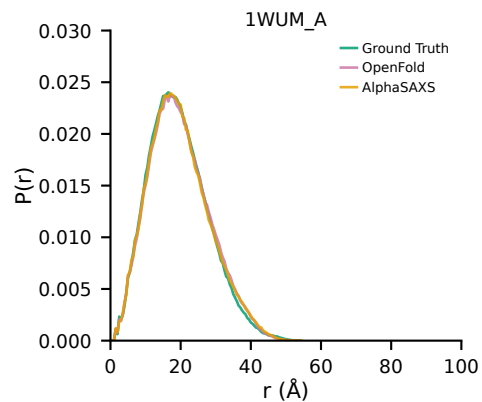

Apo

Holo

Apo

Holo

### Supplementary Figure - Page 5/20

Ground Truth

OpenFold

AlphaSAXS

P(r)

--- 1K2H-4\_A (Apo) 1ZFS-13\_B (Holo) ---

OpenFold
RMSD=6.55 Å

AlphaSAXS
RMSD=6.28 Å

Ground Truth

Apo

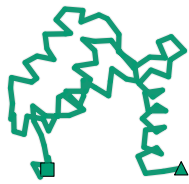

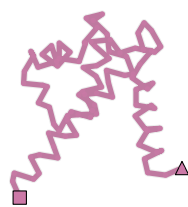

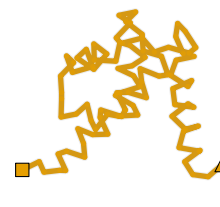

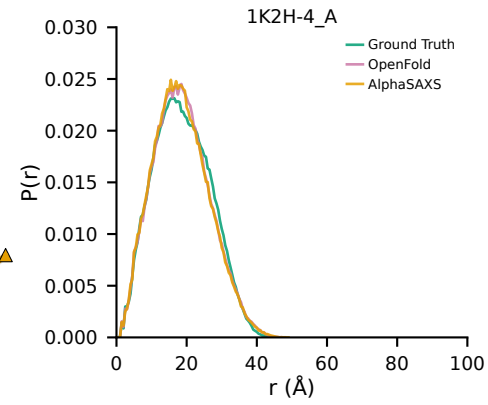

Ground Truth

OpenFold
RMSD=2.11 Å

AlphaSAXS
RMSD=2.56 Å

Holo

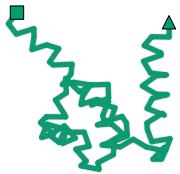

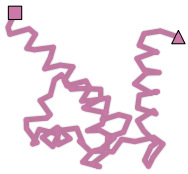

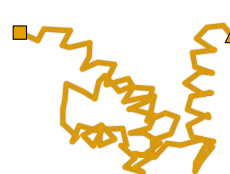

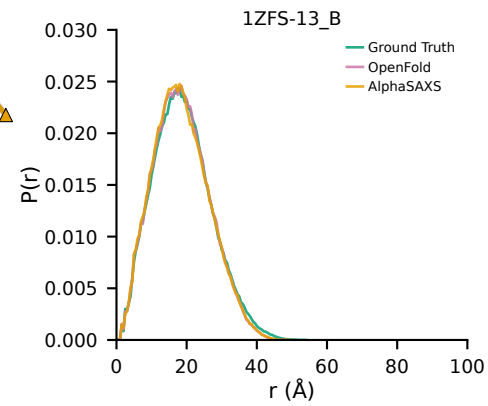

Ground Truth

--- 1LIP-2\_A (Apo) ---
OpenFold
RMSD=2.29 Å

1JTB-6\_A (Holo) ---
AlphaSAXS
RMSD=1.87 Å

Apo

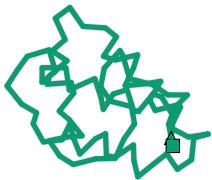

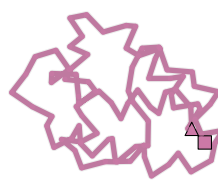

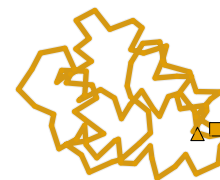

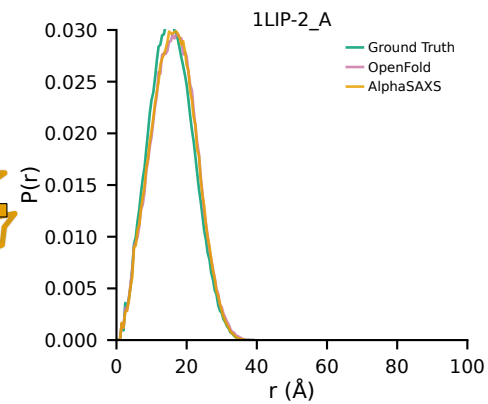

Ground Truth

OpenFold
RMSD=2.94 Å

AlphaSAXS
RMSD=2.68 Å

Holo

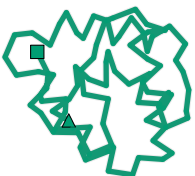

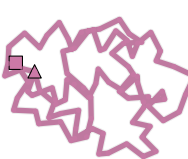

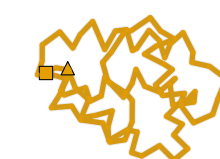

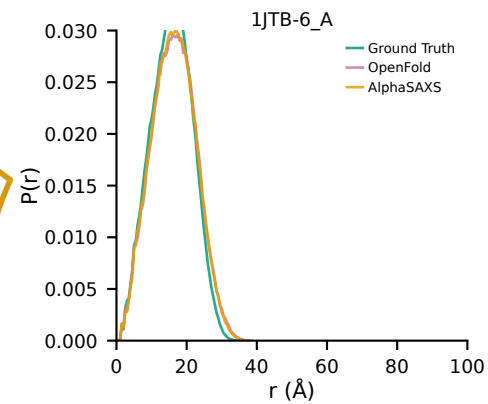

### Supplementary Figure - Page 6/20

Ground Truth

OpenFold

AlphaSAXS

P(r)

--- 1LMZ-9\_A (Apo) 1P7M-18\_A (Holo) ---

OpenFold
RMSD=3.40 Å

AlphaSAXS
RMSD=2.83 Å

Ground Truth

OpenFold

AlphaSAXS

RMSD=3.40 Å

RMSD=2.83 Å

1LMZ-9\_A

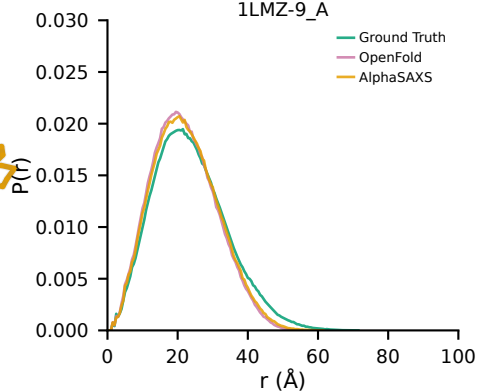

Ground Truth

OpenFold
RMSD=2.28 Å

AlphaSAXS
RMSD=2.40 Å

1P7M-18\_A

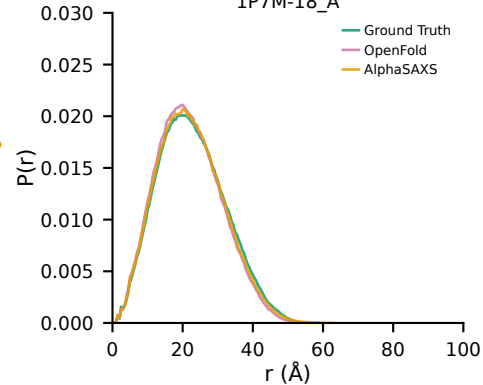

Ground Truth

--- 1M07-3\_A (Apo) ---
OpenFold
RMSD=20.12 Å

1M08-14\_A (Holo) ---
AlphaSAXS
RMSD=19.08 Å

1M07-3\_A

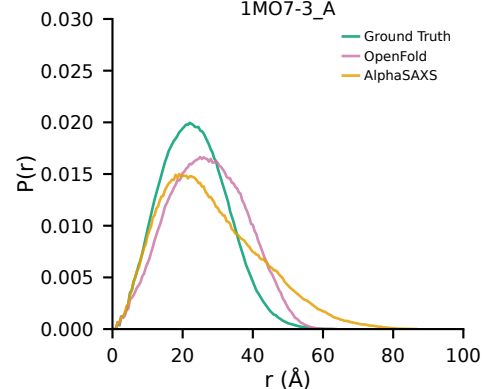

Ground Truth

OpenFold
RMSD=20.02 Å

AlphaSAXS
RMSD=18.23 Å

1M08-14\_A

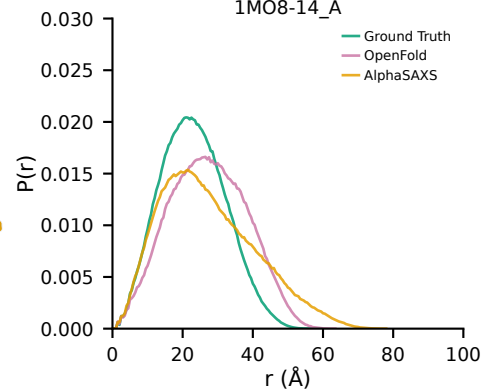

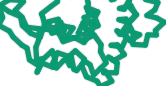

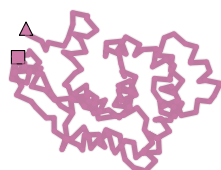

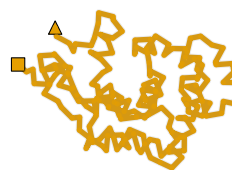

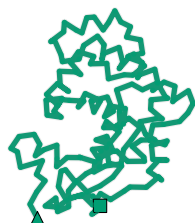

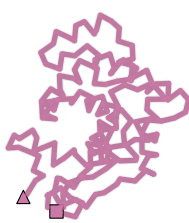

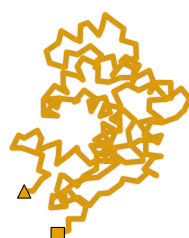

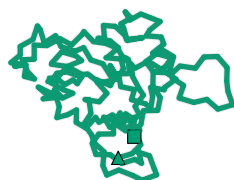

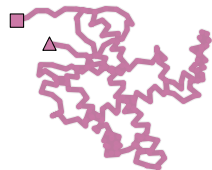

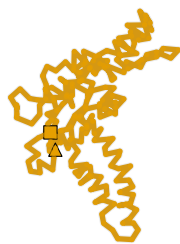

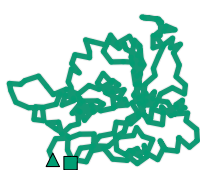

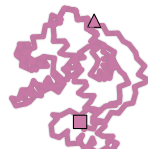

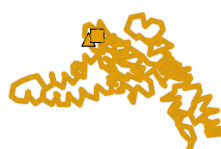

### Supplementary Figure - Page 7/20

Ground Truth

OpenFold

AlphaSAXS

P(r)

--- 1MUT-11\_A (Apo) 1PUN-7\_A (Holo) ---

OpenFold
RMSD=3.70 Å

AlphaSAXS
RMSD=3.91 Å

Ground Truth

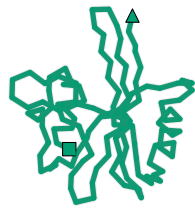

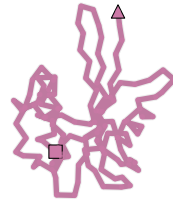

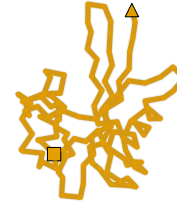

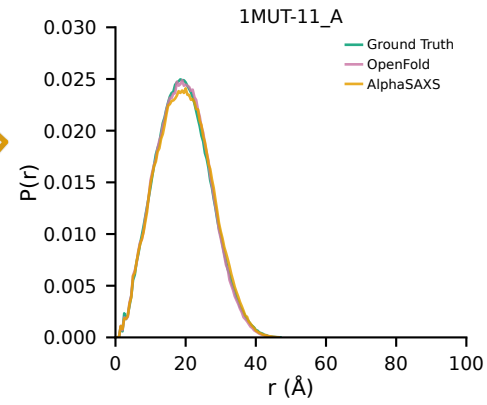

Ground Truth

OpenFold
RMSD=4.03 Å

AlphaSAXS
RMSD=3.83 Å

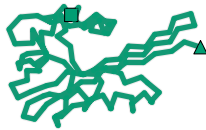

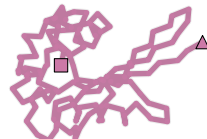

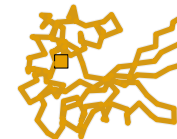

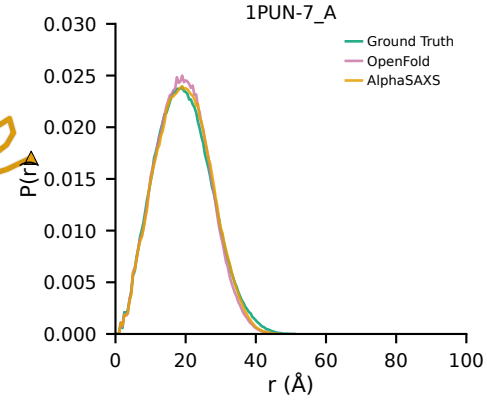

Ground Truth

--- 1INTR-7\_A (Apo) 1KRX-4\_A (Holo) ---
OpenFold
RMSD=4.37 Å

AlphaSAXS
RMSD=4.77 Å

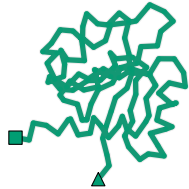

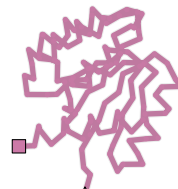

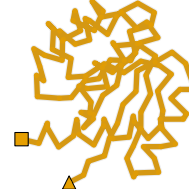

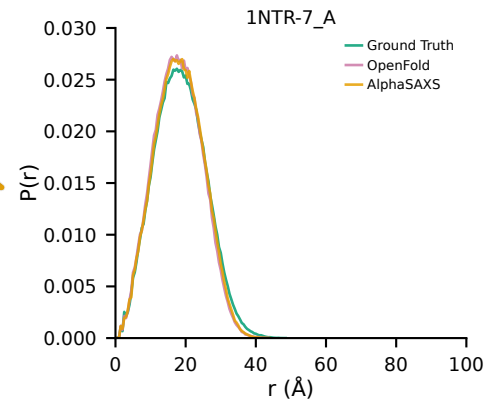

Ground Truth

OpenFold
RMSD=3.51 Å

AlphaSAXS
RMSD=3.44 Å

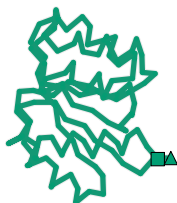

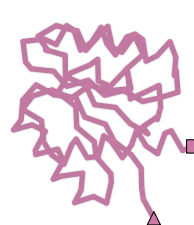

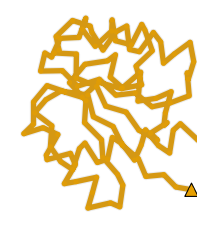

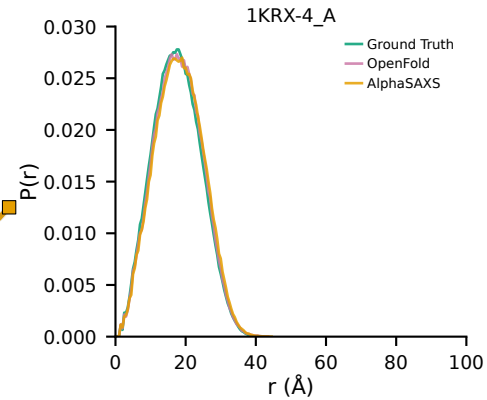

### Supplementary Figure - Page 8/20

Ground Truth

OpenFold

AlphaSAXS

P(r)

--- 1ORM-9\_A (Apo) 1QJ8\_A (Holo) ---

OpenFold
RMSD=5.65 Å

AlphaSAXS
RMSD=5.59 Å

Ground Truth

Apo

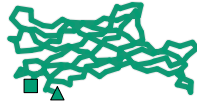

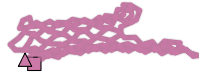

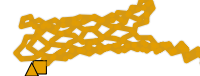

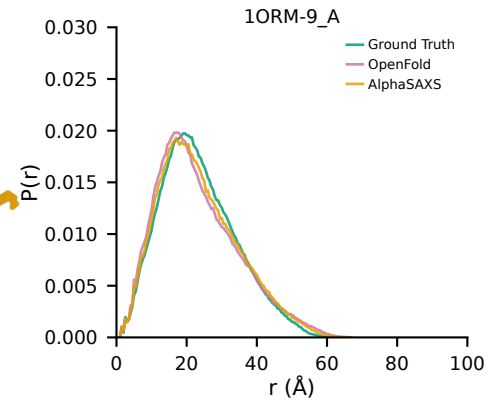

Ground Truth

OpenFold
RMSD=1.28 Å

AlphaSAXS
RMSD=2.86 Å

Holo

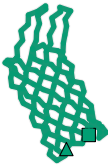

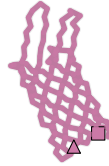

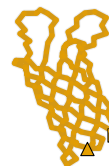

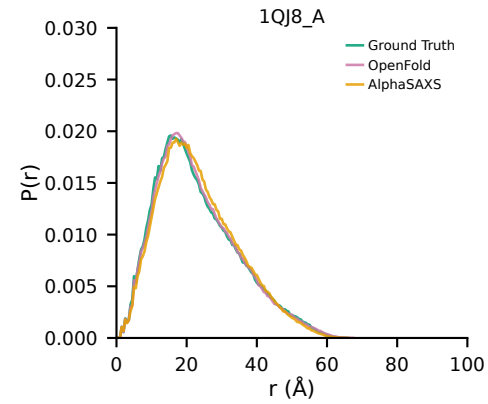

Ground Truth

--- 1S2O\_A (Apo) 1TJ5\_A (Holo) ---
OpenFold
RMSD=3.11 Å

AlphaSAXS
RMSD=2.19 Å

Apo

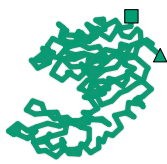

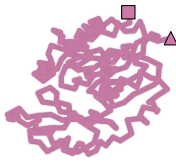

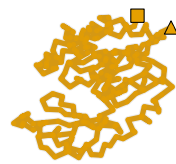

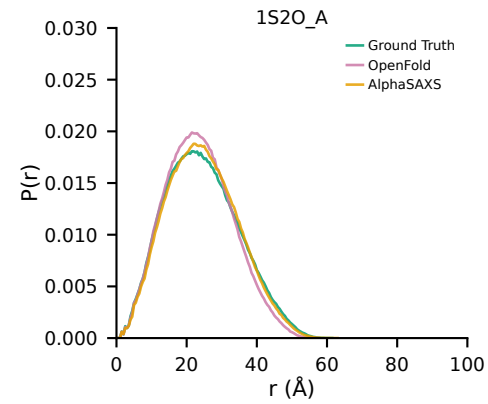

Ground Truth

OpenFold
RMSD=0.59 Å

AlphaSAXS
RMSD=1.85 Å

Holo

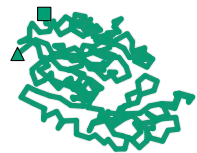

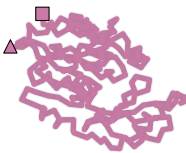

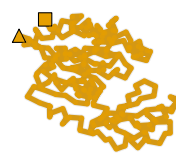

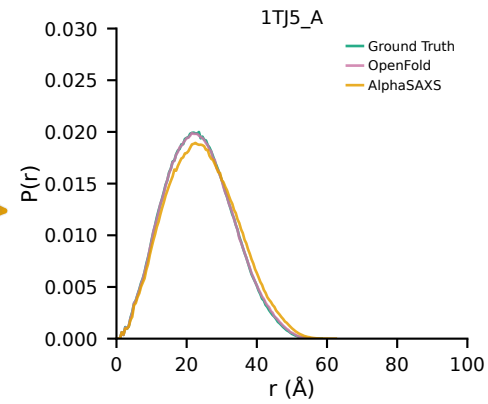

### Supplementary Figure - Page 9/20

Ground Truth

OpenFold

AlphaSAXS

P(r)

--- 1SKT-10\_A (Apo) 1TNQ-33\_A (Holo) ---

OpenFold
RMSD=5.04 Å

AlphaSAXS
RMSD=3.70 Å

Ground Truth

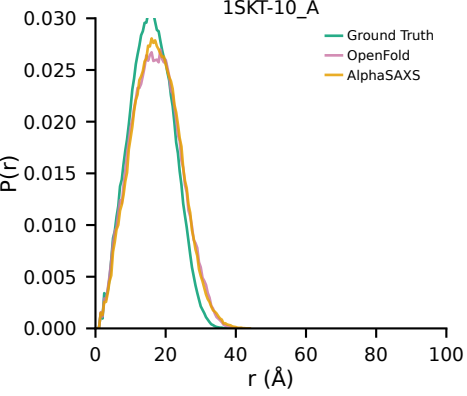

Ground Truth

OpenFold
RMSD=3.18 Å

AlphaSAXS
RMSD=5.42 Å

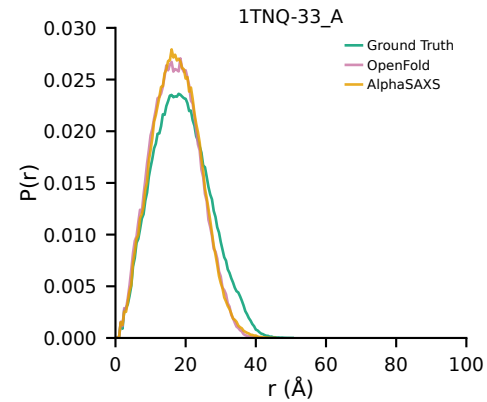

Ground Truth

--- 1SYM-2\_B (Apo) 1XYD-15\_B (Holo) ---
OpenFold
RMSD=5.83 Å

AlphaSAXS
RMSD=5.66 Å

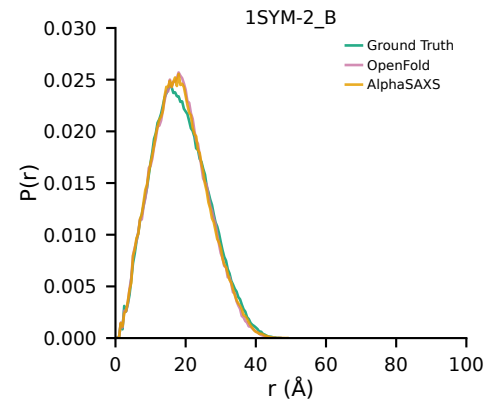

Ground Truth

OpenFold
RMSD=1.91 Å

AlphaSAXS
RMSD=2.07 Å

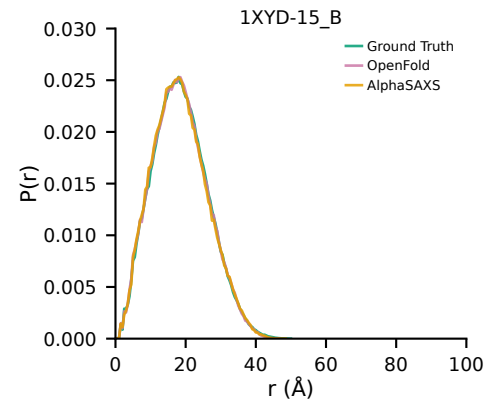

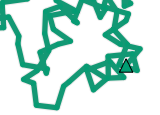

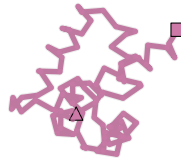

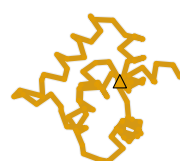

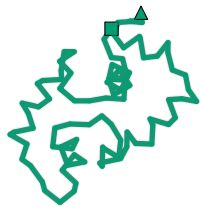

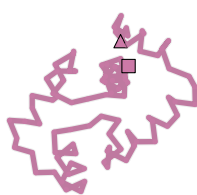

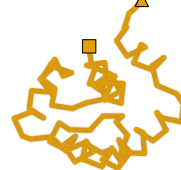

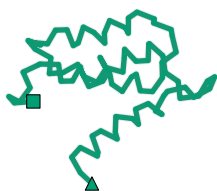

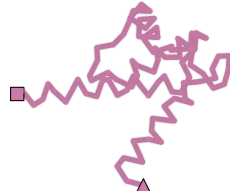

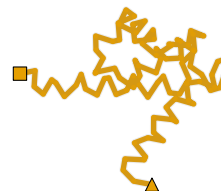

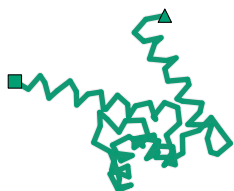

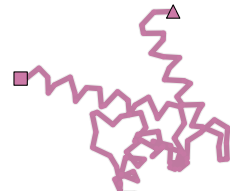

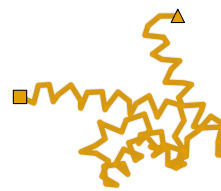

### Supplementary Figure - Page 10/20

Ground Truth

OpenFold

AlphaSAXS

P(r)

--- 1TJD\_A (Apo) 1EEJ\_B (Holo) ---
OpenFold RMSD=2.51 Å AlphaSAXS RMSD=2.14 Å

Ground Truth

OpenFold
RMSD=2.51 Å

AlphaSAXS
RMSD=2.14 Å

1TJD\_A

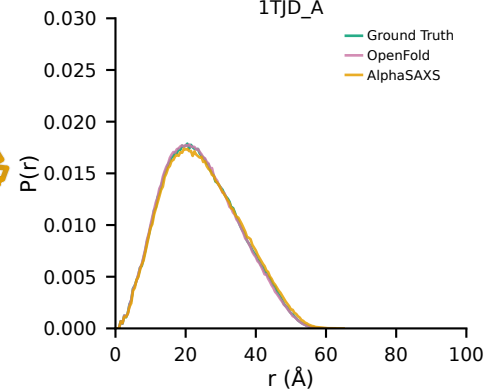

Apo

Ground Truth

OpenFold
RMSD=3.02 Å

AlphaSAXS
RMSD=2.18 Å

1EEJ\_B

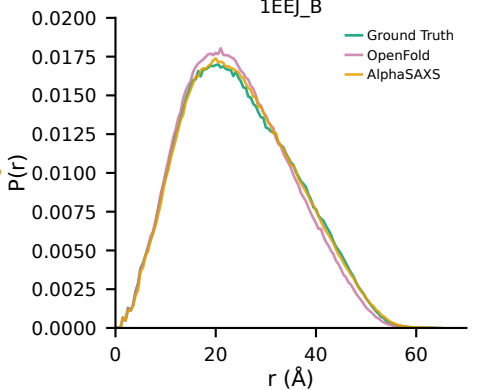

Holo

Ground Truth

--- 1VIY\_C (Apo) 1VHL\_A (Holo) ---
OpenFold RMSD=2.78 Å AlphaSAXS RMSD=2.39 Å

1VIY\_C

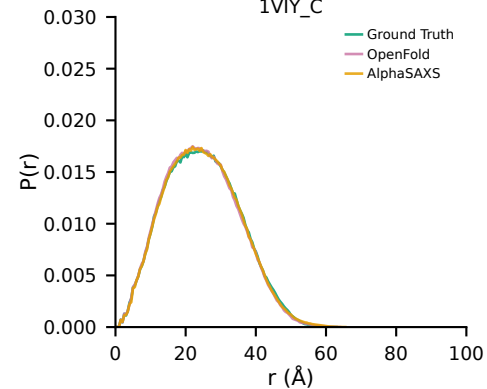

Apo

Ground Truth

OpenFold
RMSD=2.55 Å

AlphaSAXS
RMSD=2.11 Å

1VHL\_A

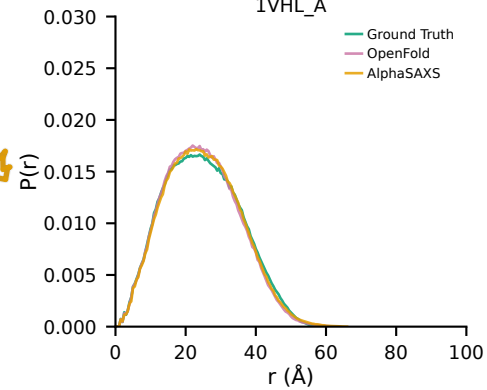

Holo

### Supplementary Figure - Page 11/20

Ground Truth

OpenFold

AlphaSAXS

P(r)

--- 1W4U-9\_A (Apo) 1UR6-4\_A (Holo) ---
OpenFold RMSD=2.27 Å AlphaSAXS RMSD=2.31 Å

Ground Truth

OpenFold
RMSD=2.17 Å

AlphaSAXS
RMSD=2.32 Å

1W4U-9\_A

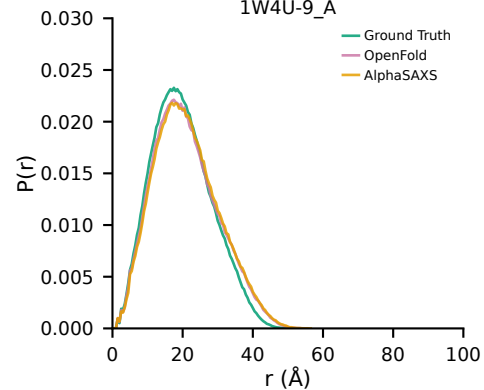

Ground Truth

OpenFold
RMSD=2.17 Å

AlphaSAXS
RMSD=2.32 Å

1UR6-4\_A

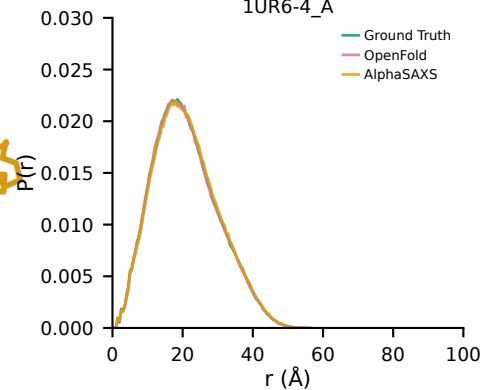

Ground Truth

--- 1WD7\_B (Apo) 1WCW\_A (Holo) ---
OpenFold RMSD=5.10 Å AlphaSAXS RMSD=4.68 Å

1WD7\_B

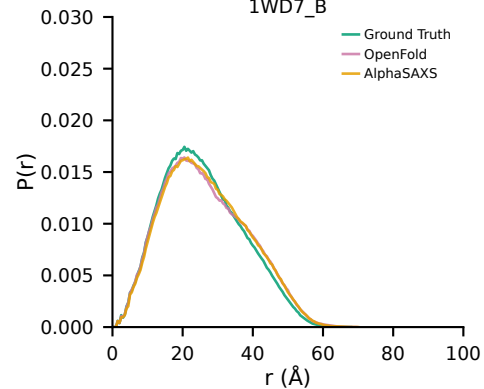

Ground Truth

OpenFold
RMSD=1.81 Å

AlphaSAXS
RMSD=1.54 Å

1WCW\_A

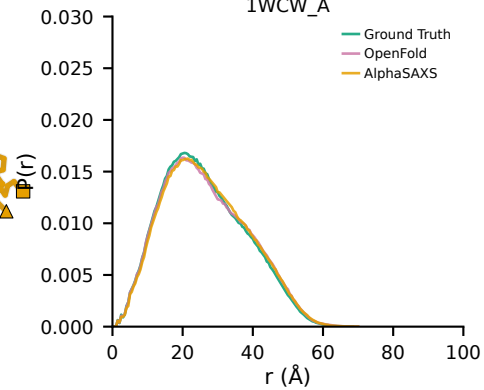

Apo

Holo

Apo

Holo

### Supplementary Figure - Page 12/20

Ground Truth

OpenFold

AlphaSAXS

P(r)

--- 1XSA-23\_A (Apo) 1XSC-20\_A (Holo) ---

OpenFold
RMSD=3.97 Å

AlphaSAXS
RMSD=3.97 Å

Ground Truth

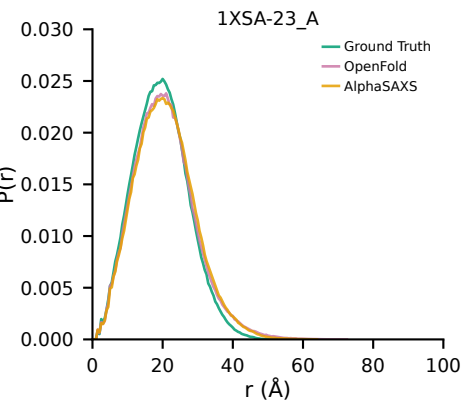

Ground Truth

OpenFold
RMSD=3.06 Å

AlphaSAXS
RMSD=2.49 Å

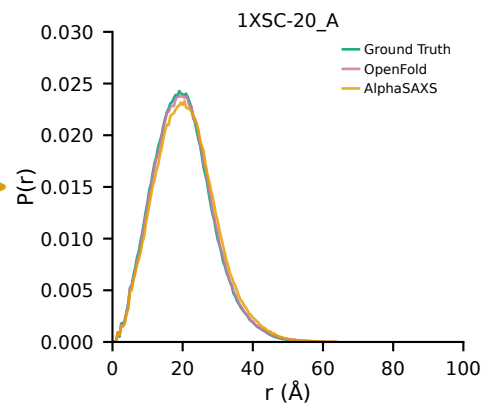

Ground Truth

--- 1ZOL\_A (Apo) ---
OpenFold
RMSD=2.85 Å

1O03\_A (Holo) ---
AlphaSAXS
RMSD=2.61 Å

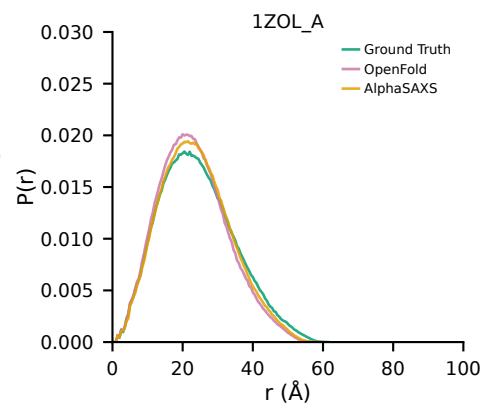

Ground Truth

OpenFold
RMSD=0.96 Å

AlphaSAXS
RMSD=2.32 Å

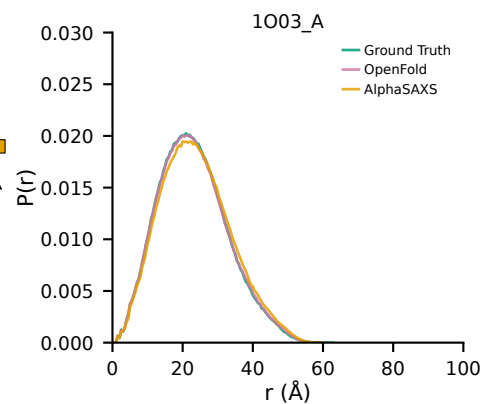

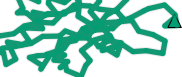

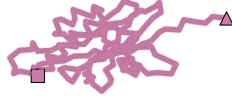

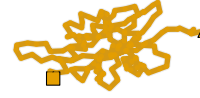

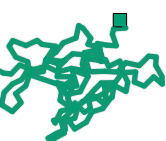

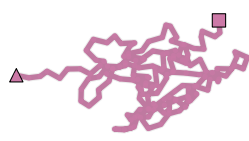

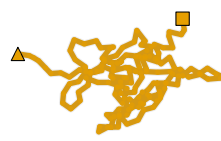

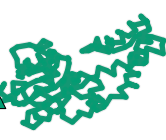

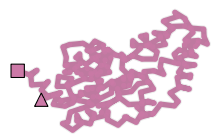

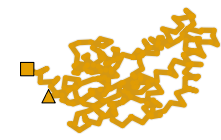

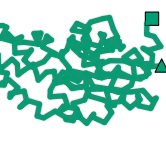

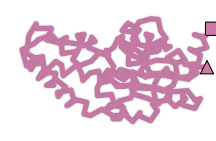

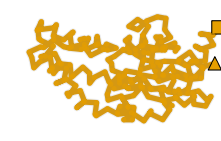

### Supplementary Figure - Page 13/20

Ground Truth

OpenFold

AlphaSAXS

P(r)

--- 2AI6-18\_A (Apo) 2OZW-3\_A (Holo) ---

OpenFold
RMSD=2.48 Å

AlphaSAXS
RMSD=2.77 Å

Ground Truth

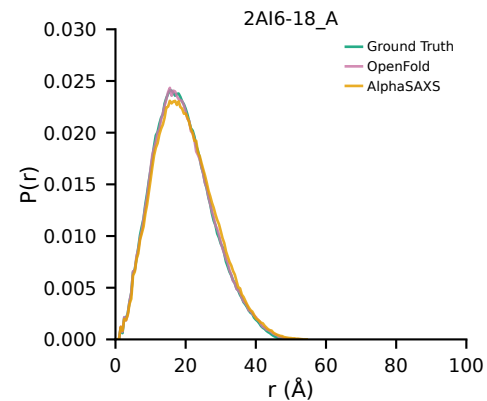

Ground Truth

OpenFold
RMSD=3.19 Å

AlphaSAXS
RMSD=3.58 Å

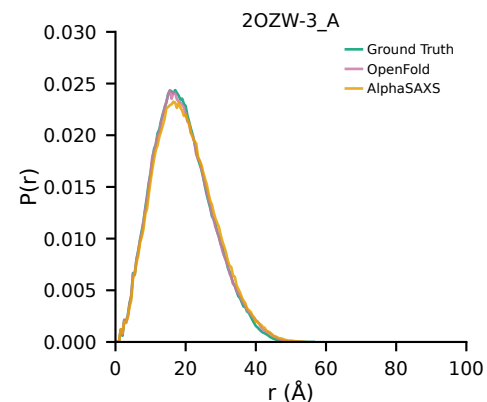

Ground Truth

--- 2CG7\_A (Apo) 2CG6\_A (Holo) ---
OpenFold
RMSD=0.93 Å

AlphaSAXS
RMSD=2.47 Å

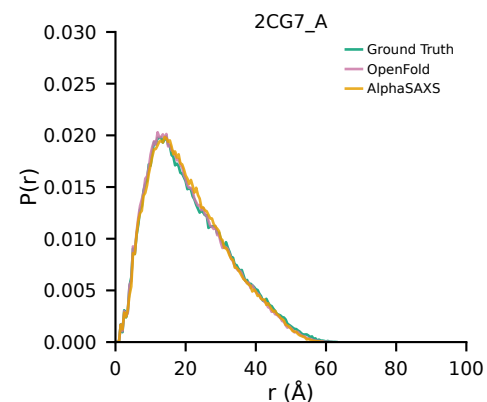

Ground Truth

OpenFold
RMSD=6.22 Å

AlphaSAXS
RMSD=7.36 Å

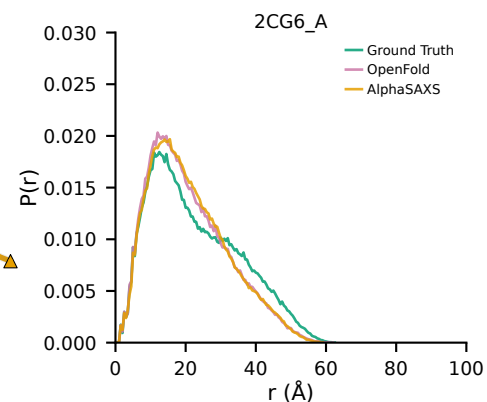

Apo

Holo

Apo

Holo

### Supplementary Figure - Page 14/20

Ground Truth

OpenFold

AlphaSAXS

P(r)

--- 2CJO-5\_A (Apo) 1ROE-10\_A (Holo) ---

OpenFold
RMSD=1.48 Å

AlphaSAXS
RMSD=1.49 Å

Ground Truth

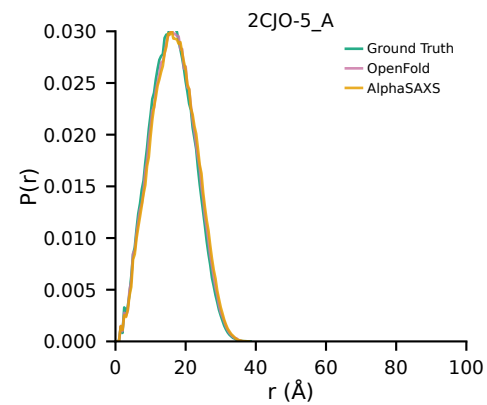

Ground Truth

OpenFold
RMSD=4.44 Å

AlphaSAXS
RMSD=4.40 Å

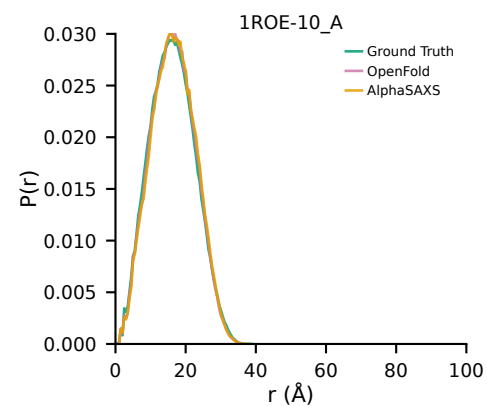

Ground Truth

--- 2D9E-12\_A (Apo) 2RS9-13\_B (Holo) ---

OpenFold
RMSD=3.84 Å

AlphaSAXS
RMSD=3.53 Å

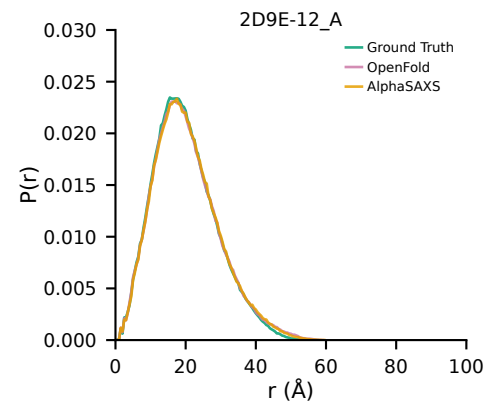

Ground Truth

OpenFold
RMSD=2.83 Å

AlphaSAXS
RMSD=3.46 Å

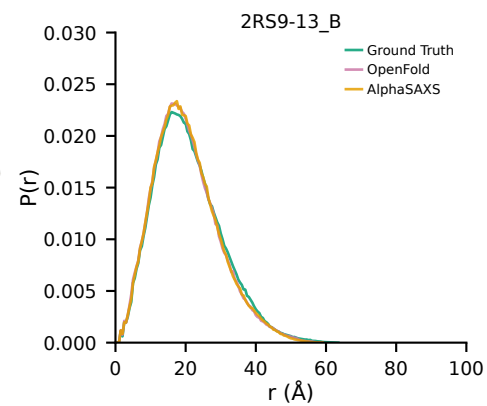

Ground Truth

### Supplementary Figure - Page 15/20

Ground Truth

OpenFold

AlphaSAXS

P(r)

--- 2F63-4\_A (Apo) 1EQM\_A (Holo) ---

OpenFold
RMSD=1.95 Å

AlphaSAXS
RMSD=2.30 Å

Ground Truth

OpenFold
RMSD=3.98 Å

AlphaSAXS
RMSD=3.44 Å

Ground Truth

--- 2JU3-3\_A (Apo) 2JU8-9\_A (Holo) ---
OpenFold
RMSD=3.03 Å

AlphaSAXS
RMSD=3.05 Å

Ground Truth

OpenFold
RMSD=1.80 Å

AlphaSAXS
RMSD=1.92 Å

Ground Truth

2F63-4\_A

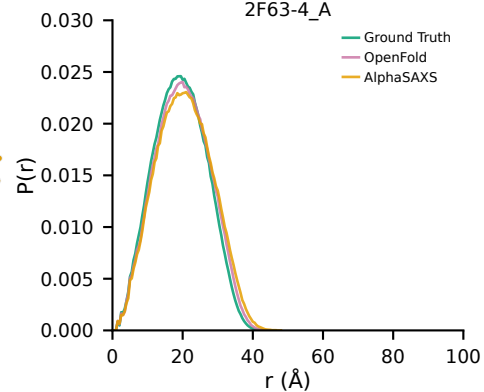

1EQM\_A

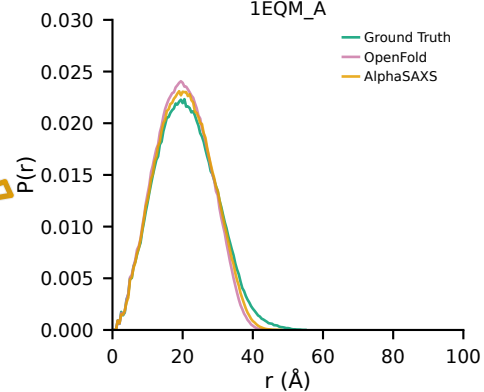

2JU3-3\_A

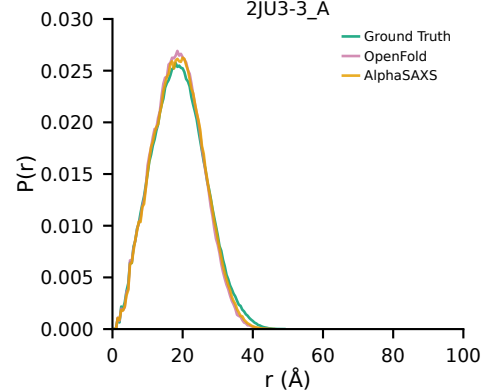

2JU8-9\_A

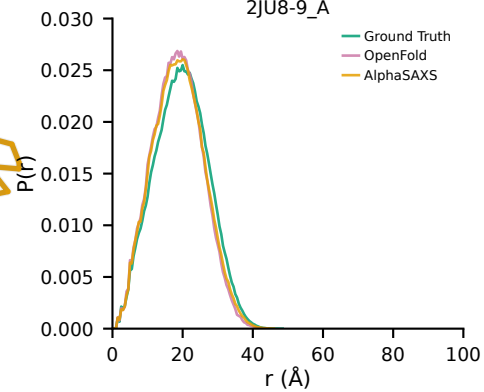

Apo

Holo

Apo

Holo

### Supplementary Figure - Page 16/20

Ground Truth

OpenFold

AlphaSAXS

P(r)

--- 2JWW-13\_A (Apo) 1RTP\_1 (Holo) ---

OpenFold
RMSD=2.66 Å

AlphaSAXS
RMSD=2.70 Å

Ground Truth

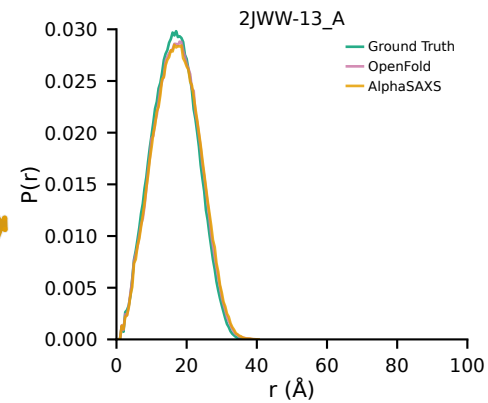

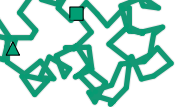

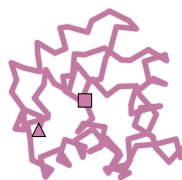

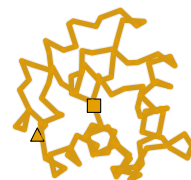

Ground Truth

OpenFold
RMSD=0.45 Å

AlphaSAXS
RMSD=1.05 Å

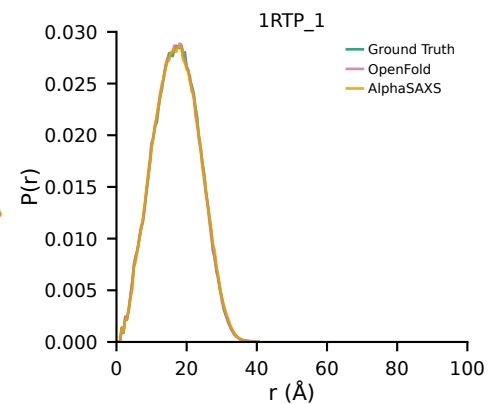

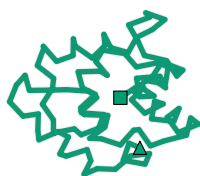

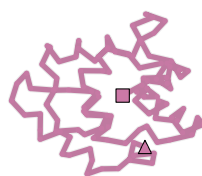

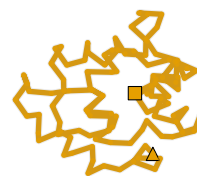

Ground Truth

--- 2K43-1\_A (Apo) 2K8R-1\_A (Holo) ---
OpenFold
RMSD=2.86 Å

AlphaSAXS
RMSD=2.71 Å

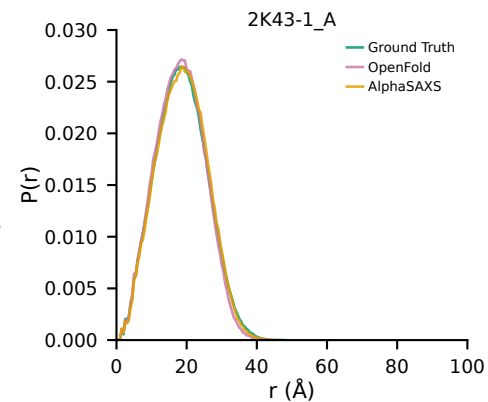

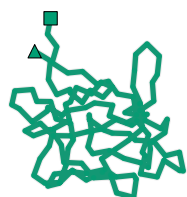

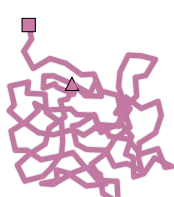

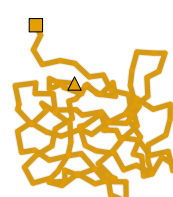

Ground Truth

OpenFold
RMSD=4.02 Å

AlphaSAXS
RMSD=3.70 Å

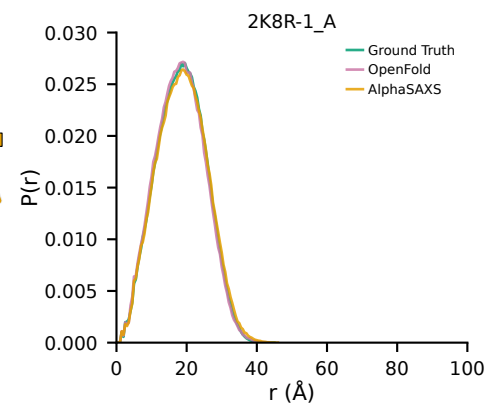

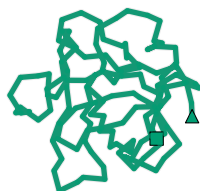

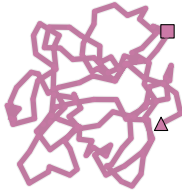

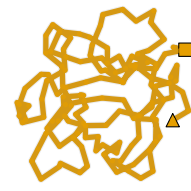

$P(r)$

AlphaSAXS
RMSD=3.76 Å

Holo

Holo

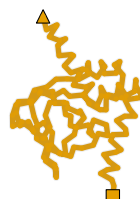

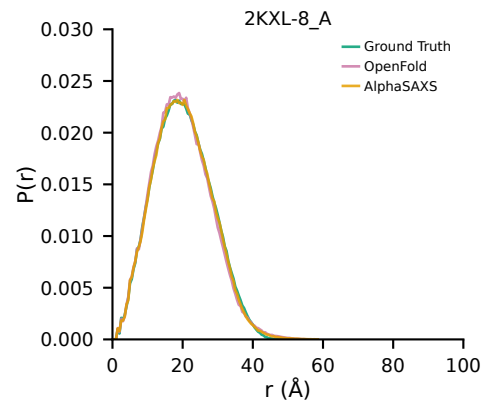

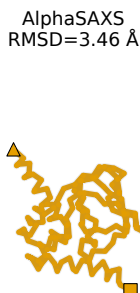

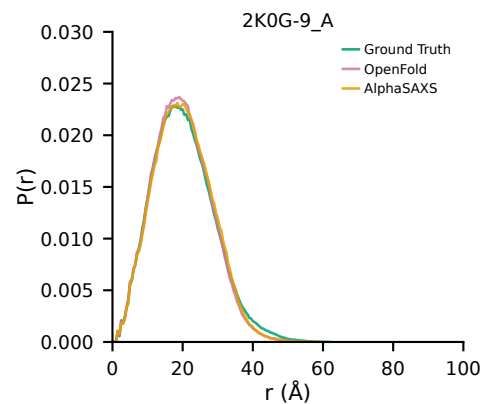

AlphaSAXS
RMSD=4.07 Å

2L50-24\_A

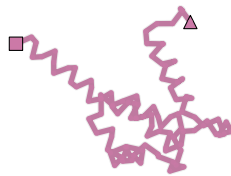

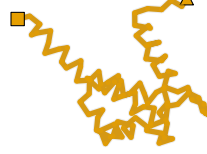

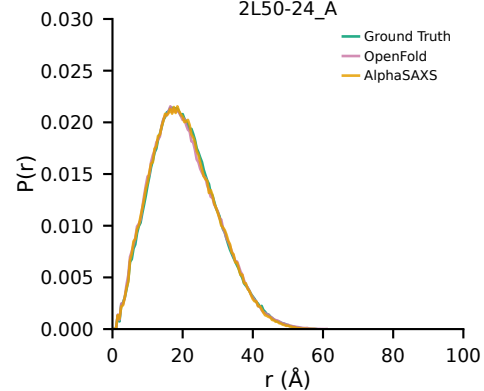

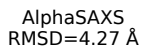

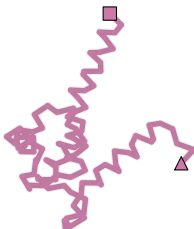

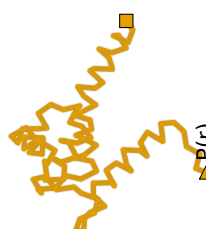

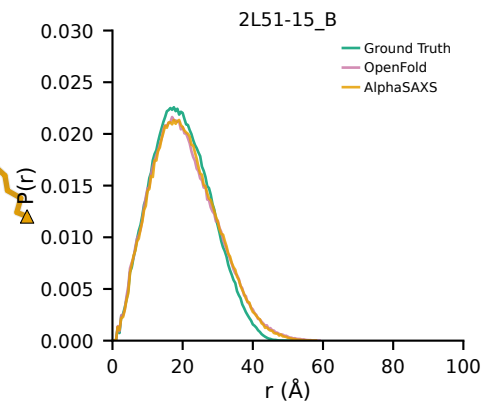

### Supplementary Figure - Page 18/20

Ground Truth

OpenFold

AlphaSAXS

P(r)

--- 2LAO\_A (Apo) 1LAH\_E (Holo) ---

OpenFold
RMSD=4.67 Å

AlphaSAXS
RMSD=3.56 Å

Ground Truth

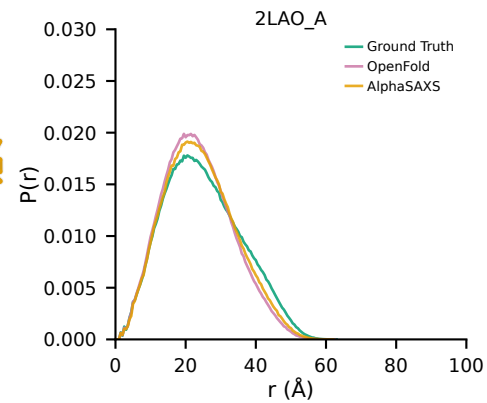

Ground Truth

OpenFold
RMSD=0.43 Å

AlphaSAXS
RMSD=1.23 Å

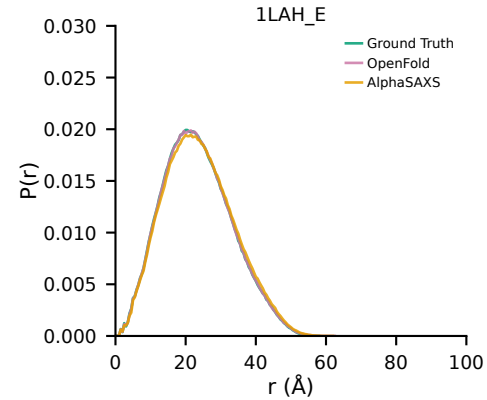

Ground Truth

--- 2LKC-4\_A (Apo) 2LKD-18\_A (Holo) ---
OpenFold
RMSD=8.13 Å

AlphaSAXS
RMSD=7.84 Å

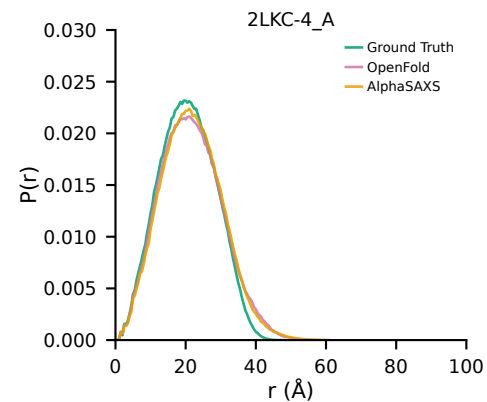

Ground Truth

OpenFold
RMSD=4.77 Å

AlphaSAXS
RMSD=5.83 Å

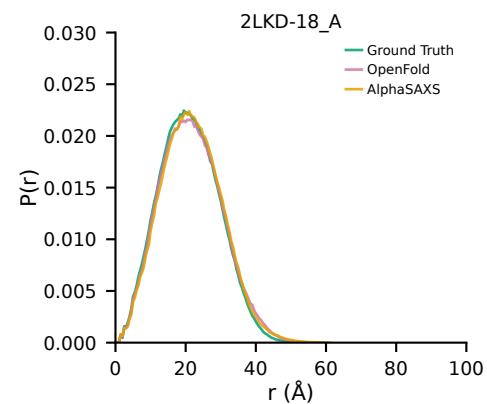

Apo

Holo

Apo

Holo

### Supplementary Figure - Page 19/20

Ground Truth

OpenFold

AlphaSAXS

P(r)

--- 2NLN-4\_A (Apo) 1RRO\_A (Holo) ---

OpenFold
RMSD=3.89 Å

AlphaSAXS
RMSD=3.82 Å

Ground Truth

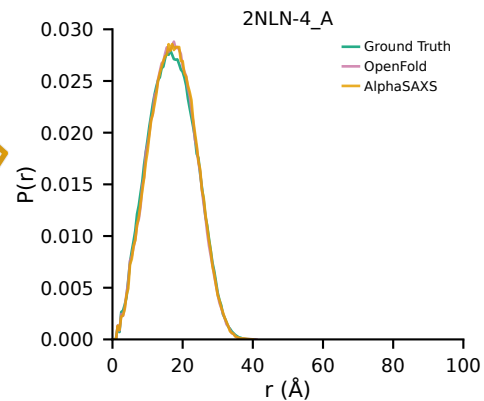

Apo

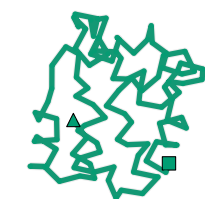

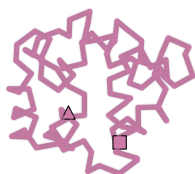

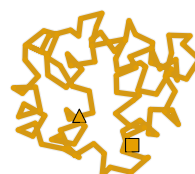

Ground Truth

OpenFold
RMSD=0.61 Å

AlphaSAXS
RMSD=1.12 Å

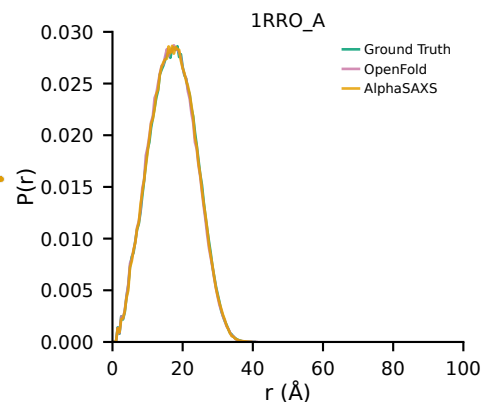

Holo

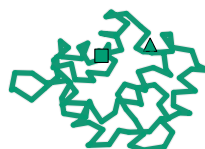

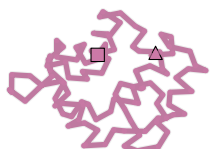

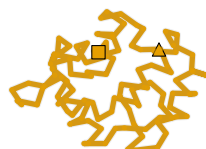

Ground Truth

--- 2RCS\_H (Apo) 1AJ7\_H (Holo) ---
OpenFold
RMSD=5.31 Å

AlphaSAXS
RMSD=4.46 Å

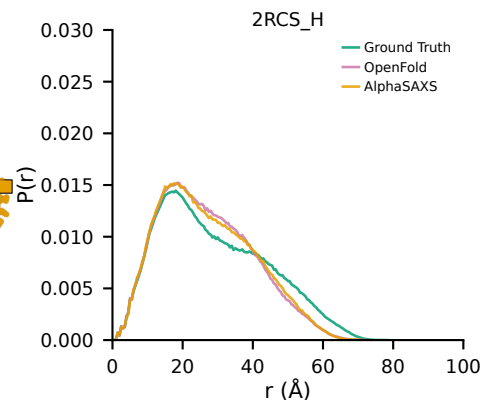

Apo

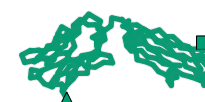

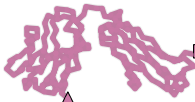

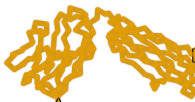

Ground Truth

OpenFold
RMSD=2.62 Å

AlphaSAXS
RMSD=4.75 Å

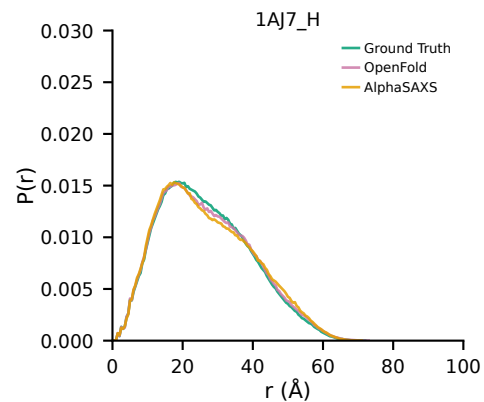

Holo

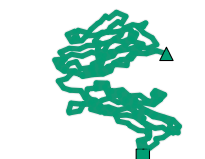

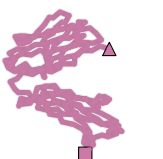

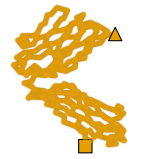

### Supplementary Figure - Page 20/20

Ground Truth

OpenFold

AlphaSAXS

P(r)

--- 2UZ5-9\_A (Apo) 2VCD-8\_A (Holo) ---

OpenFold
RMSD=3.15 Å

AlphaSAXS
RMSD=2.99 Å

Ground Truth

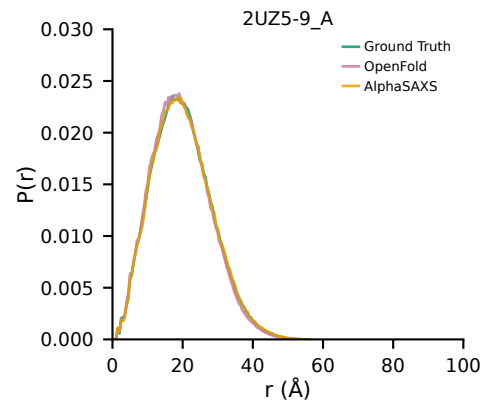

Ground Truth

OpenFold
RMSD=1.51 Å

AlphaSAXS
RMSD=1.45 Å

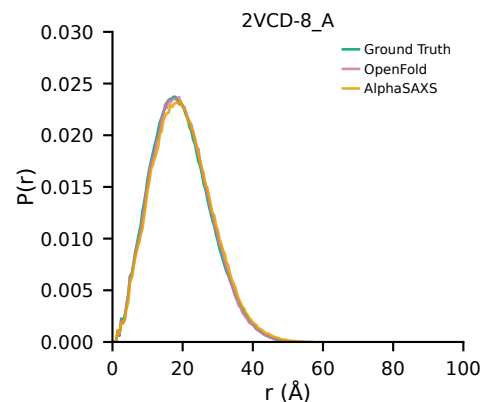

Ground Truth

--- 4AKE\_B (Apo) 2ECK\_B (Holo) ---
OpenFold
RMSD=21.21 Å

AlphaSAXS
RMSD=18.98 Å

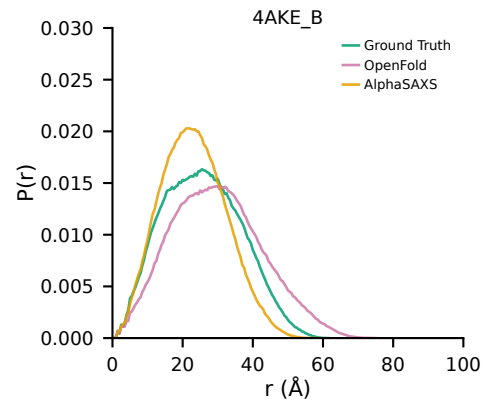

Ground Truth

OpenFold
RMSD=22.53 Å

AlphaSAXS
RMSD=18.46 Å

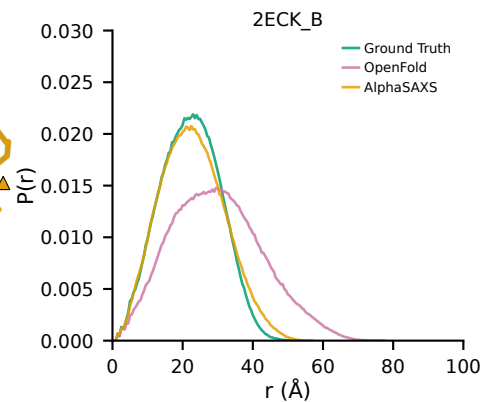

Apo

Holo

Apo

Holo
